# A comprehensive examination of Chelicerate genomes reveals no evidence for a whole genome duplication among spiders and scorpions

**DOI:** 10.1101/2024.02.05.578966

**Authors:** Gregg W.C. Thomas, Michael T.W. McKibben, Matthew W. Hahn, Michael S. Barker

## Abstract

Whole genome duplications (WGDs) can be a key event in evolution, playing a role in both adaptation and speciation. While WGDs are common throughout the history of plants, only a few examples have been proposed in metazoans. Among these, recent proposals of WGD events in Chelicerates, the group of Arthropods that includes horseshoe crabs, ticks, scorpions, and spiders, include several rounds in the history of horseshoe crabs, with an additional WGD proposed in the ancestor of spiders and scorpions. However, many of these inferences are based on evidence from only a small portion of the genome (in particular, the *Hox* gene cluster); therefore, genome-wide inferences with broader species sampling may give a clearer picture of WGDs in this clade. Here, we investigate signals of WGD in Chelicerates using whole genomes from 17 species. We employ multiple methods to look for these signals, including gene tree analysis of thousands of gene families, comparisons of synteny, and signals of divergence among within-species paralogs. We test several scenarios of WGD in Chelicerates using multiple species trees as a backbone for all hypotheses. While we do find support for at least one WGD in the ancestral horseshoe crab lineage, we find no evidence for a WGD in the history of spiders and scorpions using any genome-scale method. This study not only sheds light on genome evolution and phylogenetics within Chelicerates, but also demonstrates how a combination of comparative methods can be used to investigate signals of ancient WGDs.

## Introduction

Whole genome duplications (WGDs) occur when an individual retains both sets of chromosomes from one or more parents. While such events are often highly deleterious, occasionally the combination of novel genetic material can provide advantages that allow the whole genome duplication to propagate, resulting in a polyploid species with more than 2n chromosomes in its genome. WGDs have been important evolutionary events, with some evidence pointing to an association between environmental stress and the success of polyploid species (Van de Peer, et al. 2021). WGDs are common in plants (Masterson 1994; Adams and Wendel 2005; Barker, et al. 2016; Initiative 2019), but there are also a smaller number of important genome duplications in the history of fungi (Wolfe and Shields 1997; Ma, et al. 2009) and vertebrates (Ohno 1970; Furlong and Holland 2002; McLysaght, et al. 2002).

A common process in the evolution of polyploid species is diploidization, which is the loss of many of the excess genes and chromosomes that resulted from the WGD (Li, et al. 2021). The end result of diploidization is a return of the gene-content of the polyploid species to a nearly diploid state, with most paralogous genes that resulted from the WGD being lost or unidentifiable as paralogs (Wolfe 2001). Nevertheless, even in paleopolyploid species that have had ancient WGDs and have undergone diploidization, signatures of the WGD can remain in their genomes. For example, an excess of paralogs in the genome will have an origin that approximately coincides with the timing of the WGD. The timing of such events can be determined by multiple methods. One class of methods, generally referred to as gene tree-species tree reconciliation, uses gene tree topologies to map duplication events onto branches of the species tree (Pfeil, et al. 2005; Cannon, et al. 2015; Thomas, et al. 2017; Yan, et al. 2022). These topological methods can also potentially identify the mode of polyploidy (Thomas, et al. 2017) and can more accurately identify independent WGDs when diploidization occurs during speciation (Redmond, et al. 2023). A second class of methods examines pairwise divergence between paralogs in the same species, with the expectation that a WGD event will lead to a peak of synonymous divergence (*K*_S_) between paralogs (Lynch and Conery 2000; Blanc and Wolfe 2004; Tiley, et al. 2018). Finally, there may also be syntenic evidence for the WGD in polyploids, where whole paralogous regions of the same genome (including both coding and non-coding sequence) trace their history to the WGD event (Tang, et al. 2008; Hao, et al. 2021).

Recently, WGDs have been proposed in the history of the Arthropod sub-phylum Chelicerata, which includes horseshoe crabs, sea spiders, mites, ticks, scorpions, and spiders. In horseshoe crabs, counts of gene duplications, paralog divergence estimates, and syntenic blocks all suggest that a whole genome duplication has occurred during their evolution (Nossa, et al. 2014; Shingate, Ravi, Prasad, Tay, et al. 2020). Examination of the *Hox* gene cluster has also been used to suggest that there have been anywhere between one and three WGDs during the course of horseshoe crab evolution (Kenny, et al. 2016; Shingate, Ravi, Prasad, Tay, et al. 2020; Shingate, Ravi, Prasad, Tay and Venkatesh 2020). Similar approaches also form the basis for the claim that a WGD has occurred in the lineage ancestral to extant spiders and scorpions (Sharma, et al. 2014; Clarke, et al. 2015; Schwager, et al. 2017; Leite, et al. 2018; Fan, et al. 2021; Harper, et al. 2021; Aase-Remedios, et al. 2023). In both cases, the number of genes or genomes used for analysis has been limited. In addition, while the duplication of a conserved gene cluster (i.e. the *Hox* cluster) may be indicative of a larger (perhaps whole genome) duplication event, it is too limited a dataset with which to confirm such an event. As well as issues with the amount of data used for inferences, recent evidence supports an alternate placement of horseshoe crabs in the chelicerate phylogeny. Traditionally, the aquatic horseshoe crabs have been thought to be sister to all arachnids (spiders, scorpions, mites, and ticks), which are mostly terrestrial (Weygoldt and Paulus 1979). However, the possibility of polyphyletic origins of arachnids has been considered (see Shultz 1990) and some molecular studies have supported a scenario of polyphyletic arachnids (Sharma, et al. 2014; Ballesteros and Sharma 2019; Ontano, et al. 2021). Recently, Ballesteros, et al. (2022) presented strong evidence for horseshoe crabs being nested within arachnids, sister to spiders and scorpions, making arachnids polyphyletic. This newly proposed species tree could substantially impact how WGDs are inferred within this group when phylogenetic methods are used (McKibben, et al. 2024).

Here, we use whole-genome sequences from 17 chelicerate species, in combination with several different analytical methods, to look for ancient WGDs in this group. These methods include gene tree reconciliation, synonymous divergence between paralogs, and whole-genome analyses of synteny. Using multiple species trees as a backbone for analysis, we find no evidence for a WGD taking place in the history of spiders and scorpions. In contrast, our suite of methods all find some evidence for at least one WGD occurring during the evolution of horseshoe crabs, even in light of their new placement in the chelicerate phylogeny.

## Methods

### Data

To investigate the possible existence of whole genome duplication (WGD) events in chelicerates on a genome-wide scale, we took a multi-faceted approach. We initially downloaded 18 chelicerate genomes with annotations available at the beginning of this project from various sources: NCBI’s Assembly database (https://www.ncbi.nlm.nih.gov/assembly) Ensembl Metazoa (Yates, et al. 2022; release 51), the i5k database (Consortium 2013; Thomas, et al. 2020), and, for two samples, the data supplements of their genome publications (Fan, et al. 2021; Nong, et al. 2021). These genomes span the various taxonomic groups contained within the subphylum Chelicerata, including four species from the superorder Parasitiformes (mites and ticks), two species from the superorder Acariformes (mites), eight species from the order Araneae (spiders), one species from the order Scorpiones (scorpions), and four species from the order Xiphosura (horseshoe crabs) (Fig. 1). For this study, we treat Parasitiformes and Acariformes as orders. For phylogenetic analyses, we also include two insects (*Drosophila melanogaster* and *Bombyx mori*) as outgroups for tree rooting. See Supplemental Table S1 for full details of the samples and summaries of their assemblies and annotations.

**Figure 1:**
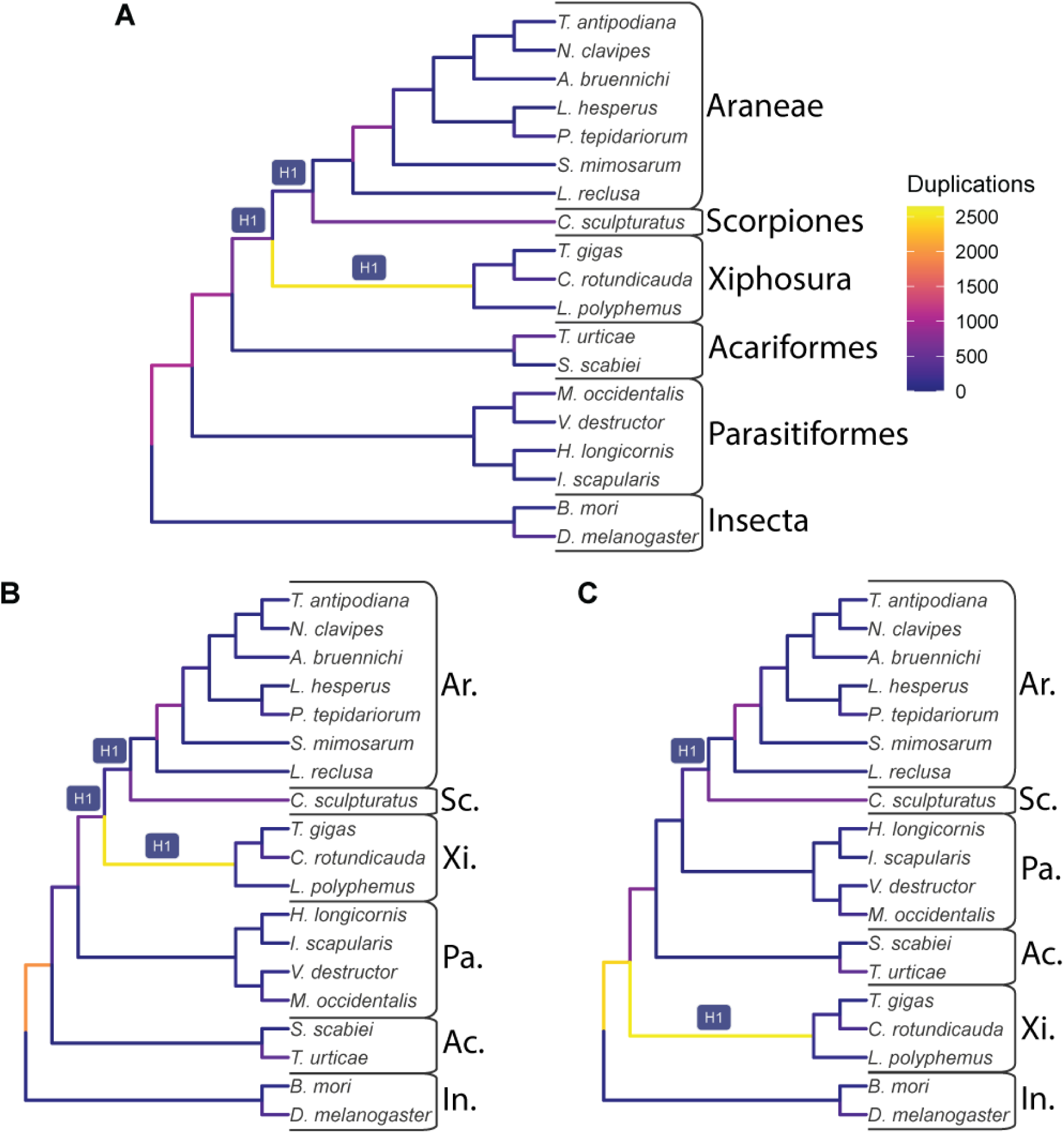
The input species trees used with GRAMPA, which are also the lowest scoring trees when considering possible WGDs at the branches labeled H1. Branches are shaded by the number of duplications that map to them. A) The species tree topology inferred in this study from 11,016 gene families. B) The species tree inferred by Ballesteros, et al. (2022). C) A species tree that places horseshoe crabs (Xiphosura) sister to Arachnids. For all B and C, taxonomic groups are labeled as follows: Ar. = Araneae (spiders); Sc. = Scorpiones (scorpions); Xi. = Xiphosura (horseshoe crabs); Ac. = Acariformes (mites); Pa. = Parasitiformes (mites and ticks); In. = Insecta (insects).

We observed that annotations of one of the horseshoe crabs, *Tachypleus tridentatus*, contained 79,557 genes, more than twice as many as any other species in our sample, including the other horseshoe crabs. While on the surface this may indeed be indicative of a recent WGD in this species, we also note that the median gene length for this species is only 1,377 bp. While this is not the shortest gene length in our sample, it is considerably smaller than the rest of the horseshoe crabs, which all have a median gene length of over 8,500 bp (see Supplemental Table S1). Because this could be indicative of annotation error in this species and because we are interested in ancient rather than recent WGDs, we excluded this sample from our analyses. In total, our final dataset contained 17 chelicerate species and 2 outgroup insects for analyses that span almost 600 million years of genome evolution.

### Gene tree reconciliation analysis

We extracted the coding sequence of the longest transcript from each gene in each of our 19 species and used FastOrtho (https://github.com/olsonanl/FastOrtho), which is a reimplementation of orthomcl (Li, et al. 2003), to cluster genes into gene families. Using an inflation value of 3, we inferred 49,561 gene families. We then extracted the sequences in each gene family, correcting for inconsistencies resulting from the data originating from various sources and aligned each gene family with Guidance2 (Sela, et al. 2015) using MAFFT (Katoh and Standley 2013) as the underlying aligner, and removing any alignment columns with a score below 0.93. We also performed our own alignment filtering by removing columns in sliding windows of 3 codons that have 2 codons with 2 or more gaps in 50% of the sequences in that alignment. We also removed any sequences that were made up of greater than 20% gap characters and removed any alignments with sequences from fewer than 4 species or that were shorter than 33 codons after all filtering. See Supplementary Table S2 for alignment filtering details.

We translated the remaining 11,016 alignments from nucleotides to amino acids and inferred gene trees with IQ-TREE (Nguyen, et al. 2015) using ultrafast bootstrap (Hoang, et al. 2018); the gene trees were used to infer a species tree with ASTRAL-Multi (Rabiee, et al. 2019). For subsequent reconciliation analyses, we rooted our gene and species trees using the outgroup insects with Newick Utilities (nw_reroot; Junier and Zdobnov 2010). Gene trees that could not be rooted because there was no outgroup were excluded from reconciliation analyses. After rooting, we retained gene trees from 6,368 gene families. To further reduce possible gene tree inference error, we used bootstrap rearrangement implemented in Notung (Chen, et al. 2000) with a bootstrap threshold of 90. This method forces inferred duplications on branches in our gene trees with a bootstrap score less than this threshold to be resolved in such a way that minimizes the number of duplications and losses counted in the tree.

We used these 6,368 rooted, bootstrap-resolved gene trees and a species tree as input to GRAMPA (Thomas, et al. 2017) to identify the placement of any WGDs in the chelicerate phylogeny. Briefly, GRAMPA performs least common ancestor (LCA) mapping from each gene tree to the species tree but allows for WGDs to be present in the species tree by representing them as multi-labeled trees (MUL-trees), in which one or more tip labels appear twice. By comparing LCA mapping scores between the input species tree and a set of MUL-trees defined by target lineages, GRAMPA can determine if a WGD has occurred on a hypothesized lineage. For our runs, we set as target lineages for WGD identification those on which WGDs have previously been proposed: specifically, the branch leading to spiders and scorpions and the branch leading to horseshoe crabs. We also used multiple different species trees as input to GRAMPA to test the same scenarios. In addition to the species tree we inferred using ASTRAL (Fig. 1A), the two alternate species tree topologies we tested were a recently inferred phylogeny from Ballesteros, et al. (2022)—in which horseshoe crabs group within arachnids, specifically sister to spiders and scorpions (Fig. 1B)— and a ‘traditional’ species tree topology, in which horseshoe crabs are sister to all arachnid species (Fig. 1C). For the ‘traditional’ tree, because of the unresolved placement of Acariformes and Parasitiformes (Sharma, et al. 2014; Ontano, et al. 2021), we simply use the topology recovered by Ballesteros, et al. (2022) and manually placed horseshoe crabs sister to arachnids.

### Synteny analysis

We used estimates of synteny to test for paleopolyploid ancestry in each of our 19 species. Self-self syntenic analyses for each genome were made using MCScanX (Wang, et al. 2012). We used the default settings of MCScanX to detect and visualize intraspecific syntenic blocks. Given that ancient WGDs may be highly fractionated, we also used a minimum block size of 3 to recover potentially highly fragmented blocks of synteny.

### Synonymous divergence between paralogs (K**_S_**)

To construct gene families and to estimate the age distribution of gene duplications we used the DupPipe pipeline (Barker, et al. 2008; Barker, et al. 2010). Briefly, DupPipe translates coding transcripts from nucleotide to peptide sequences and identifies reading frames by comparing Genewise (Birney, et al. 2004) alignments to the best-hit protein from a collection of proteins from the 19 sampled genomes. For all DupPipe runs, we used protein-guided DNA alignments to align our nucleic acid sequences while maintaining the reading frame. We estimated synonymous divergence (*K*_S_) using PAML (Yang 2007) with the F3X4 model for each node in the gene-family phylogenies. We identified peaks of gene duplication as evidence for potential ancient WGDs in histograms of the age distribution of gene duplications (*K*_S_ plots). To infer ancient WGDs in the paralog age distributions we used a recently developed machine learning approach, SLEDGe (Sutherland, et al. 2024), to classify *K*_S_ plots with peaks consistent with an ancient WGD. Specifically, we applied the support vector machine classifier from SLEDGe on node *K*_S_-values for species that had greater than 1,500 gene duplicates, subsampling down to 3,000 duplicates when more than 3,000 were present. For each *K*_S_ distribution that SLEDGe predicted as being indicative of a WGD, we also used mixture modeling and manual curation to identify significant peaks of gene duplication consistent with a WGD and to estimate their median paralog *K*_S_ values. We ran normalmixEM for a maximum of 400 iterations to fit the maximum number of *k*-components for each *K*_S_ distribution selected from a likelihood ratio test available in the boot.comp function from the mixtools R library (Benaglia, et al. 2009). Finally, to assess if WGD peaks in the paralog *K*_S_ distributions were shared between species, we used OrthoPipe from EvoPipes (Barker, et al. 2008; Barker, et al. 2010) to identify orthologs between species and PAML (Yang 2007) to estimate their *K*_S_ values using the same procedure and protein database as described for the DupPipe analyses. We then assessed species divergence by estimating the median *K*_S_ of all orthologs with a *K*_S_ of 5 or lower for each species pair and compared to the median *K*_S_ of each WGD peak.

## Results

### Inference of the species tree

We used the genomes of 17 chelicerates and 2 insect outgroups to reconstruct the Chelicerata phylogeny, with an emphasis on Arachnids and horseshoe crabs. Using 11,016 gene trees we confirm the placement of Xiphosura (horseshoe crabs) as nested within Arachnids (Fig. 1A), in agreement with Ballesteros et al. (Fig 1B; Ballesteros, et al. 2022). However, our inferred tree differs from theirs in the placement of the superorders Acariformes and Parasitiformes. Our results show that Acariformes is sister to the spider, scorpion, and horseshoe crab clade, while Ballesteros, et al. (2022) suggest that Parasitiformes is more closely related to them. However, the placement of these groups is also ambiguous in their analyses and has been contentious in previous studies (Sharma, et al. 2014; Ontano, et al. 2021).

### Reconciliation analysis

We used the inferred species tree, as well as two other hypothesized sets of relationships, to test various hypotheses of WGD in the history of chelicerate evolution. Specifically, based on synteny and duplication of some gene families, multiple rounds of WGD have been proposed in horseshoe crabs (Nossa, et al. 2014; Kenny, et al. 2016; Shingate, Ravi, Prasad, Tay, et al. 2020; Shingate, Ravi, Prasad, Tay and Venkatesh 2020), and, based on the duplication of the *Hox* gene cluster, one WGD has been proposed in the ancestor of spiders and scorpions (Schwager, et al. 2017). Using gene tree topologies from thousands of genes, GRAMPA (Thomas, et al. 2017) finds no evidence for a WGD in the history of spiders and scorpions using either our inferred species tree, the Ballesteros, et al. (2022) species tree, or the traditional species tree in which horseshoe crabs are sister to Arachnids (Figs. 1 and 2). In each case, we tested whether the species tree with a WGD proposed on any of the target lineages (H1 lineages in Fig. 1) better explains the duplication history of the genes in these genomes than a species tree with no proposed WGDs. However, in each case we find that the species tree without any proposed WGDs results in the lowest duplication and loss score (black shapes in Fig. 2). Our evidence is definitive for any WGD in the history of spiders and scorpions; however, we do see evidence for a large number of duplications on the branch leading to horseshoe crabs regardless of the species tree used (Fig. 1). We also find that the second- and third-lowest scoring scenarios when using our inferred species tree posit a WGD in horseshoe crabs (Fig. 2, Supplemental Table S3, Fig. S1). The horseshoe crab clade is also often inferred as being involved in a WGD in the next lowest scoring MUL-trees when using the other two species trees, but usually in more complicated scenarios (Figs. S1 and S2; Supplemental Tables S4 and S5). That is, while GRAMPA did not find a WGD in the history of horseshoe crabs as the single most parsimonious reconciliation, there are multiple pieces of evidence that point to one or more possibly occurring.

**Figure 2:**
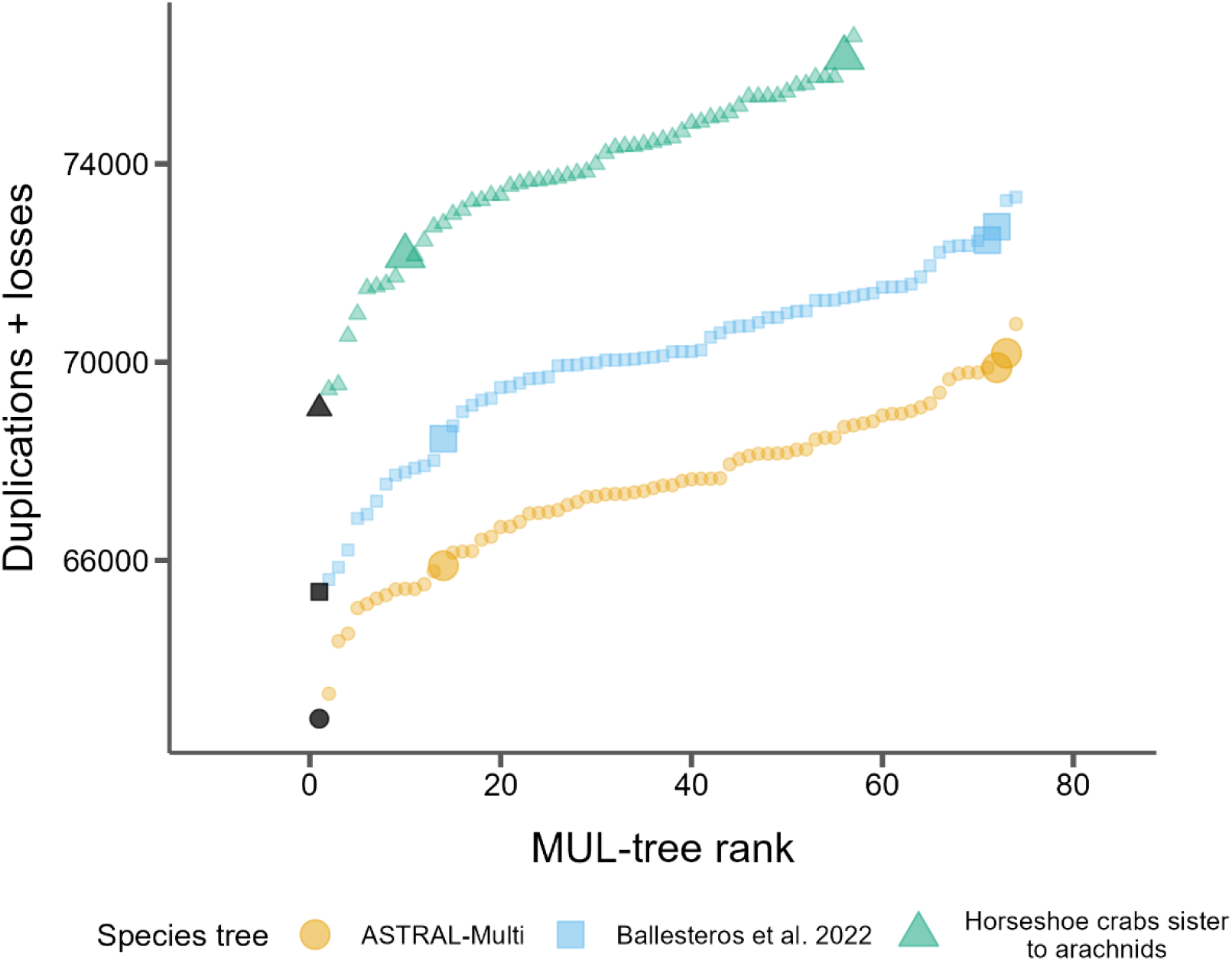
GRAMPA scores (duplications + losses) for every MUL-tree considered for each of the three species trees. Black points represent the input singly-labeled species tree with no WGD proposed. All other shaded points propose one WGD on one of the target H1 branches (see Fig. 1). Larger points indicate autopolyploidy scenarios and smaller dots indicate allopolyploidy scenarios.

We also find that, when comparing reconciliation scores between species trees, our species tree and the Ballesteros, et al. (2022) species tree both explain the history of gene duplication and loss better than the ‘traditional’ species tree in which horseshoe crabs are not nested within Arachnids (Fig. 2). This is further evidence in favor of the placement of this group as sister to spiders and scorpions. While our species tree always better explains the data from rooted gene trees than Ballesteros et al. (2002), this should not be surprising since we inferred our tree from a superset of these data (both rooted and unrooted gene trees).

### Synteny and K_S_ analyses

We next looked at other genome-wide signatures of WGDs among chelicerates. Specifically, we looked for intraspecific synteny blocks, which should be widespread in genomes that have undergone WGD, and distributions of synonymous divergence (*K*_S_) of paralogs within each genome. If a WGD has occurred in the history of a genome, a secondary peak of *K*_S_ should be present in these distributions. Across both analyses, we again find no evidence for WGD in any spider or scorpion genomes but do find suggestive evidence for at least one occurring in the history of horseshoe crabs (Fig. 3). Only two species, *C. rotundicauda* and *T. gigas*, both horseshoe crabs, showed substantial amounts of intraspecific synteny. Both of these species, along with the other horseshoe crab, *L. polyphemus*, were also predicted by SLEDGe to have signatures of WGD in their *K*_S_ distributions (Fig. 3, Supplemental Table S6). Mixture models placed the median *K*_S_ of this duplication at ∼0.85-1.35 (Fig. 3, Supplemental Table S6). The average ortholog divergence between the three horseshoe crabs was ∼0.22, compared to the average divergence with *C. sculpturatus* at ∼4.09, suggesting the WGD peak corresponds to the same branch identified with an excess number of gene duplications and losses in our gene tree topology reconciliation analysis above (Fig. 1, Fig. 3, Supplemental Table S7). In addition, one mite species, *Tetranychus urticae*, was predicted by SLEDGe to contain a WGD in its *K*_S_ distribution (Fig. 3). However, this species had few intraspecific syntenic blocks (Fig. 3; Supplemental Table S6) and no signal of excess duplication in the reconciliation analysis (Fig. 1). It is likely that the prediction made by SLEDGe in *T. urticae* is an artefact of assembly or annotation in this species.

**Figure 3:**
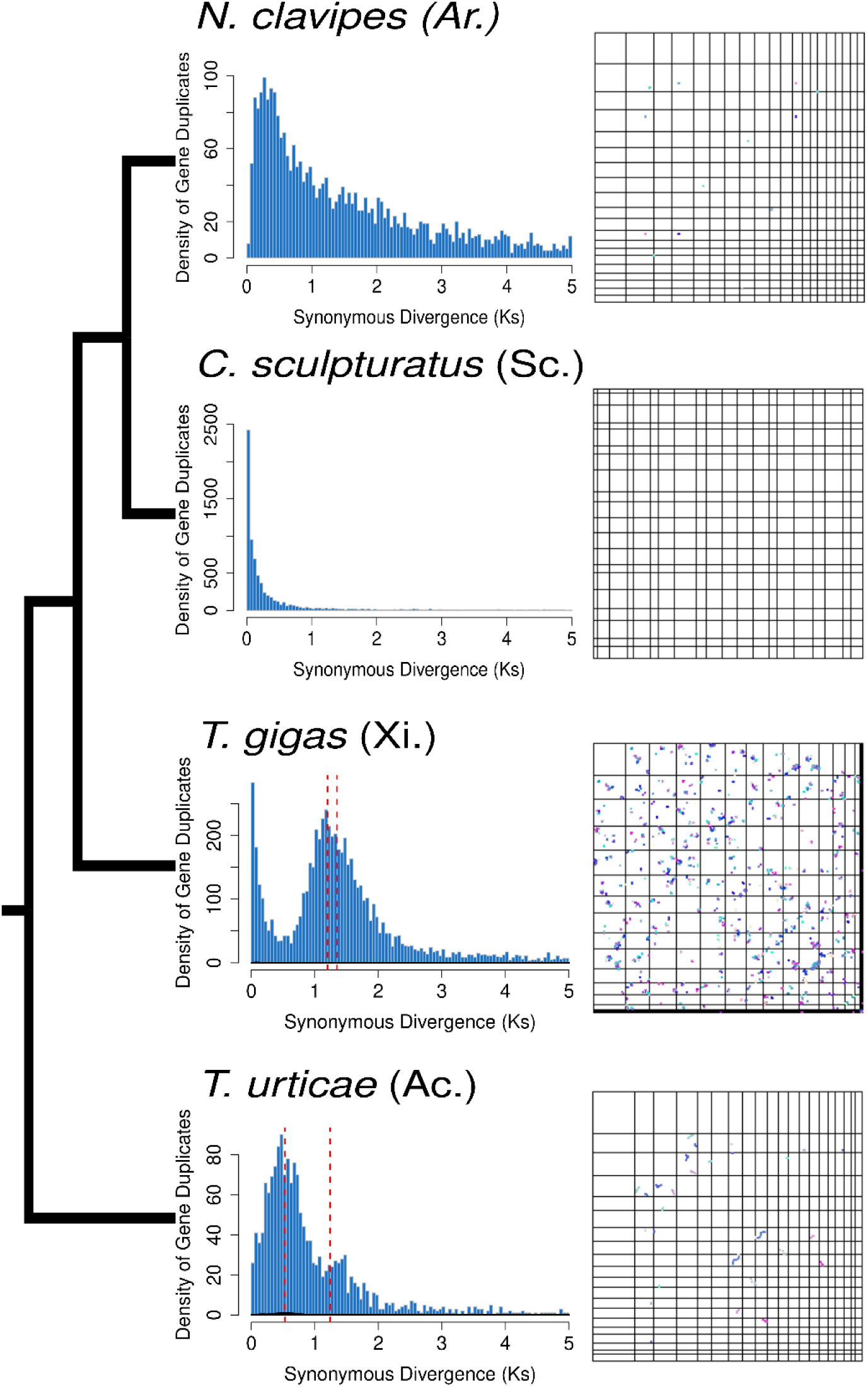
Distributions of *K*_S_ (left) and synteny (right) for select samples (See Figs. S5 and S6 for all samples) from Acariformes (Ac.), Xiphosura (Xi.), Araneae (Ar.) and Scorpiones (Sc.). These samples all showed the highest levels of synteny among samples in each group. The species tree topology is shown on the far left. Red dotted lines indicate the median K_S_ of mixture models fit to distributions that were predicted by SLEDGe to be indicative of WGDs.

## Discussion

Whole genome duplications (WGDs) can be a key event in the evolution of a species, possibly facilitating adaptation (Ohno 1970; Werth and Windham 1991; Adams and Wendel 2005; Crow and Wagner 2006). While the process of diploidization (the return of the genome to a diploid state after WGD) can make more ancient WGDs harder to detect, multiple methods have been developed that have the potential to capture the signal of these events in extant genomes. Here, we used several of these methods to investigate the existence of ancient WGDs in the Chelicerates (Nossa, et al. 2014; Kenny, et al. 2016; Shingate, Ravi, Prasad, Tay, et al. 2020; Shingate, Ravi, Prasad, Tay and Venkatesh 2020). Several rounds of WGD have been proposed in the history of horseshoe crab evolution, and a single WGD has been proposed in the ancestor of spiders and scorpions (Sharma, et al. 2014; Clarke, et al. 2015; Schwager, et al. 2017; Leite, et al. 2018; Fan, et al. 2021; Harper, et al. 2021; Aase-Remedios, et al. 2023). The evidence for these events usually starts with the observation of the duplication of a well-conserved gene family cluster, the *Hox* genes. Further investigations of intraspecific synteny, gene tree topologies, and divergence have also been used previously, but until now have been limited to only a few genes or genomes.

Using 17 chelicerate whole genomes we find no evidence for a WGD in the history of spiders and scorpions. When reconciling gene tree topologies to a species tree that allows for the inference of WGDs, the best-scoring scenario is always the one without any WGDs, regardless of the input species tree topology used. For spiders and scorpions, we also see no excess intraspecific synteny or peaks in divergence of paralogs that would indicate a WGD. This implies that the two copies of the *Hox* gene cluster observed in some spiders and scorpions may instead be the result of a more limited duplication event. While *Hox* gene clusters are thought to be relatively slowly evolving outside of WGDs, this is not always the case (Mulhair, et al. 2023; Mulhair and Holland 2024). Therefore, inferences about WGDs should not be made from the *Hox* cluster alone (e.g. Farhat, et al. 2023).

We do find some evidence for WGDs during horseshoe crab evolution. While no MUL-trees are the single-most optimal solution in the gene tree analysis, we do find a burst of gene duplications on the branch leading to horseshoe crabs. This burst is observed regardless of the species tree considered (Fig. 1). Previously, anywhere from one to three WGDs have been proposed along the horseshoe crab lineage. In fact, if multiple WGDs occurred, this may diminish the signal for any single proposed MUL-tree. Since our tests using GRAMPA are limited to a single MUL-tree, this may in turn hinder our ability to explicitly identify any single WGD as the most parsimonious scenario. In addition to the large number of duplications on the horseshoe crab lineage, we also observe notable intraspecific synteny and peaks in divergence of paralogs (Fig. 3).

In the course of our study of WGDs in Chelicerates, we also reconstructed a species tree for our 17 species (Fig. 1A). Using our whole genome data and including paralogs in our species tree inference (cf. Smith and Hahn 2021), we find that the horseshoe crabs (Xiphosura) are nested within Arachnids, directly sister to spiders (Araneae) and scorpions (Scorpiones). This agrees with several recent molecular phylogenies of this group (Sharma, et al. 2014; Ballesteros and Sharma 2019; Ontano, et al. 2021; Ballesteros, et al. 2022), and rejects a tree suggested by the biomes in which the organisms live, where the aquatic horseshoe crabs are sister to the mostly terrestrial arachnids (Fig. 1C). In this traditional monophyletic Arachnid tree, separate WGDs would need to be proposed for both spiders/scorpions and horseshoe crabs. However, the molecular trees allow the possibility that a single WGD took place in the ancestor of spiders, scorpions, and horseshoe crabs. We also tested this scenario (Fig. 1A) and were able to rule out this possibility.

Our work shows that, even for ancient polyploids, whole genome comparative evidence can still find signals of WGDs. While the duplication of a single gene family can be a good initial clue that a WGD has occurred, as it was for metazoans (Amores, et al. 1998), whole genome evidence is still needed for a more confident inference (Furlong and Holland 2002; McLysaght, et al. 2002; Hokamp, et al. 2003; Dehal and Boore 2005). Our work shows that this is also the case for Chelicerates. In horseshoe crabs, duplications in *Hox* gene clusters coincide with synteny, peaks of synonymous divergence in intraspecific paralogs, and gene duplication reconciliation in the Chelicerate phylogeny. None of these additional pieces of evidence is present in the lineage leading to spiders and scorpions. Our work also adds to the growing body of evidence that horseshoe crabs are not sister to all arachnids as was traditionally thought, but rather are placed within the arachnid group, directly sister to spiders and scorpions.

## Data availability

The genomes used in our analyses are available from their respective databases (see Supplemental Table S1). All other data generated for this project (gene alignments, gene trees, etc.) are available on TBD. Scripts used to parse and analyze this data are available at https://github.com/gwct/spider-wgd.

## Acknowledgements

We thank Zheng Li for helpful discussions on our analyses. Gene family analysis was performed on the FASRC Cannon cluster supported by the FAS Division of Science Research Computing Group at Harvard University. M.W.H. was supported by National Science Foundation grant DEB-1936187.

## Supplemental Figure Legends

*Figure S1*

The lowest scoring MUL-trees from the GRAMPA analysis using our inferred species tree.

*Figure S2*

The lowest scoring MUL-trees from the GRAMPA analysis using the Ballesteros, et al. (2022) species tree.

*Figure S3*

The lowest scoring MUL-trees from the GRAMPA analysis using a traditional species tree with horseshoe crabs sister to arachnids.

*Figure S4*

Dot plots showing intra-species synteny for all species (19 panels, labeled with species name) with a max block size of 3.

*Figure S5*

Dot plots showing intra-species synteny for all species (19 panels, labeled with species name) with a max block size of 5.

*Figure S6*

Distributions of K_s_ between paralogs of all species (19 panels, labeled with species name). Dashed red lines indicate the median K_s_ of mixture models fit to each K_s_ distribution that was indicative of a WGD.

