## Supplementary figures and images for "A comprehensive examination of Chelicerate genomes reveals no evidence for a whole genome duplication among spiders and scorpions"

### Supplemental Figure S1

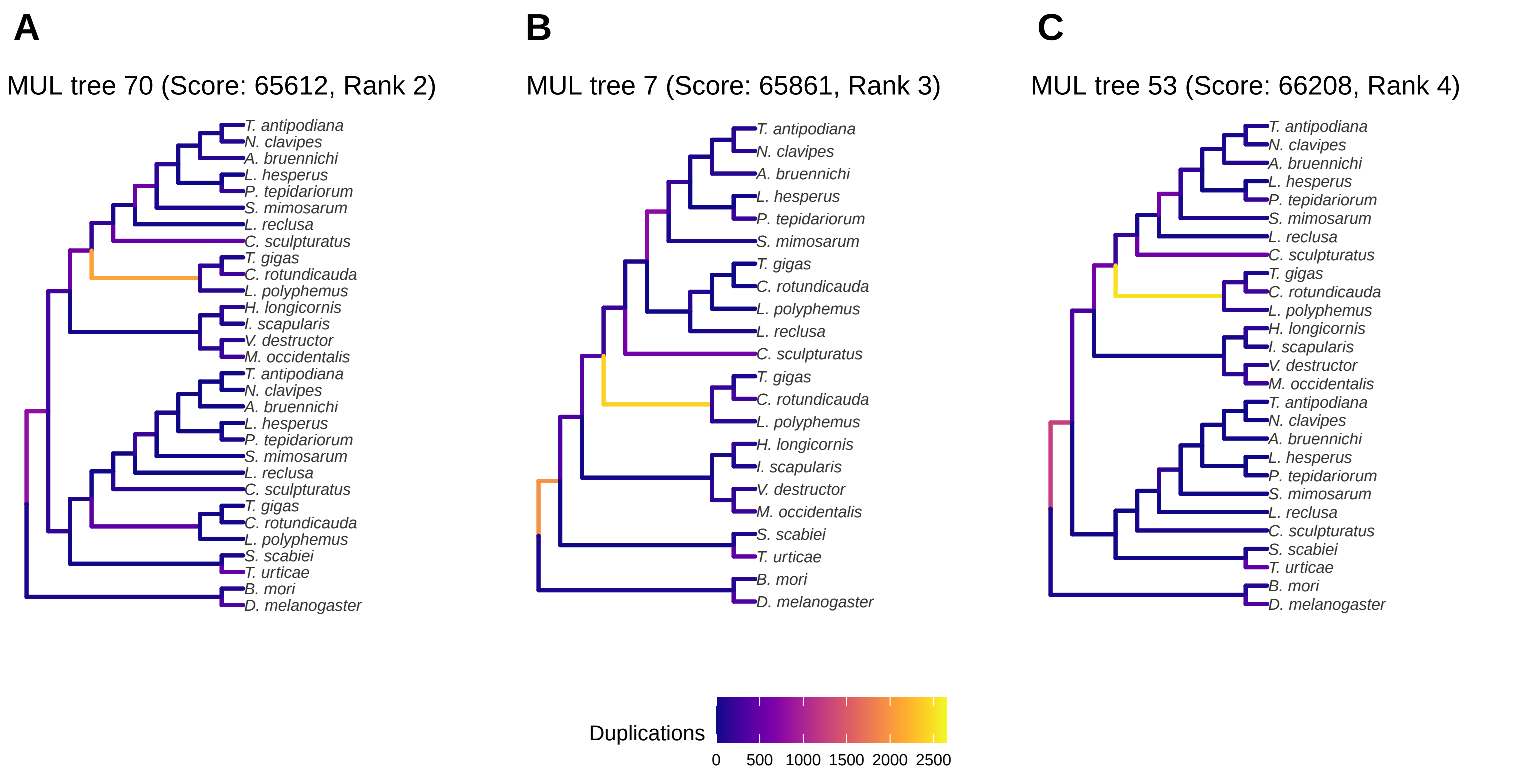

### Supplemental Figure S2

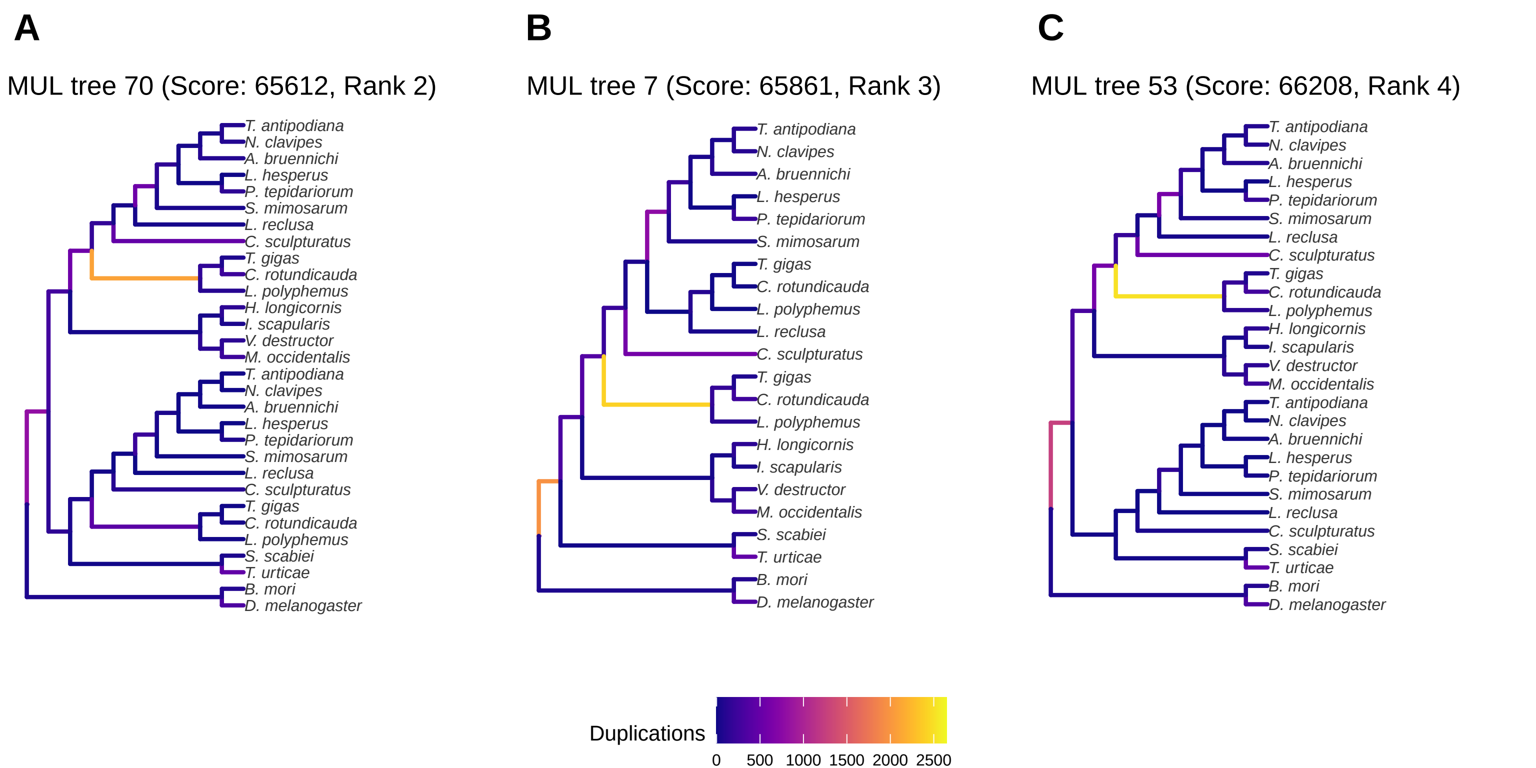

### Supplemental Figure S3

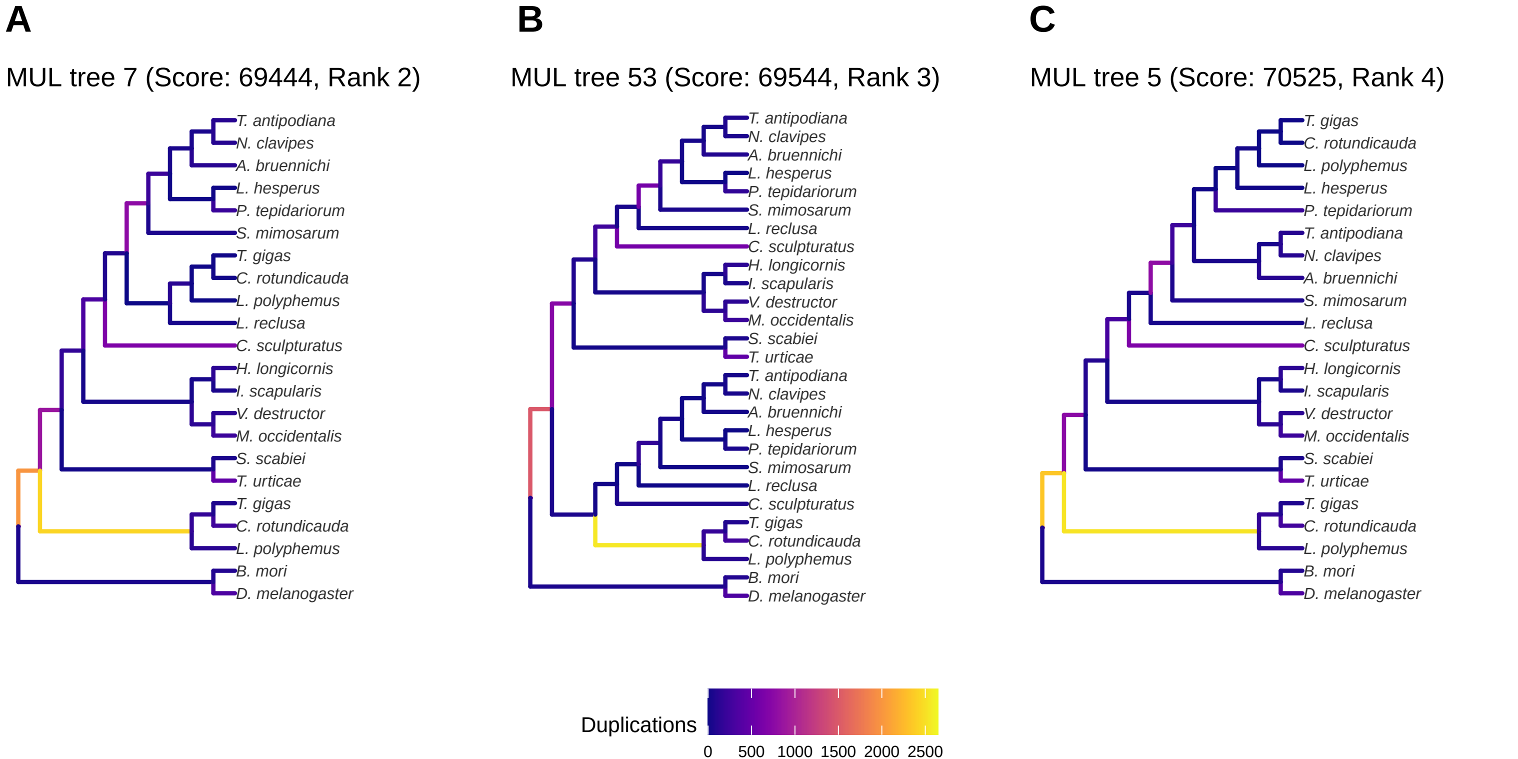
