## Supplemental Figure S4 for "A comprehensive examination of Chelicerate genomes reveals no evidence for a whole genome duplication among spiders and scorpions"

### *Argiope bruennichi*

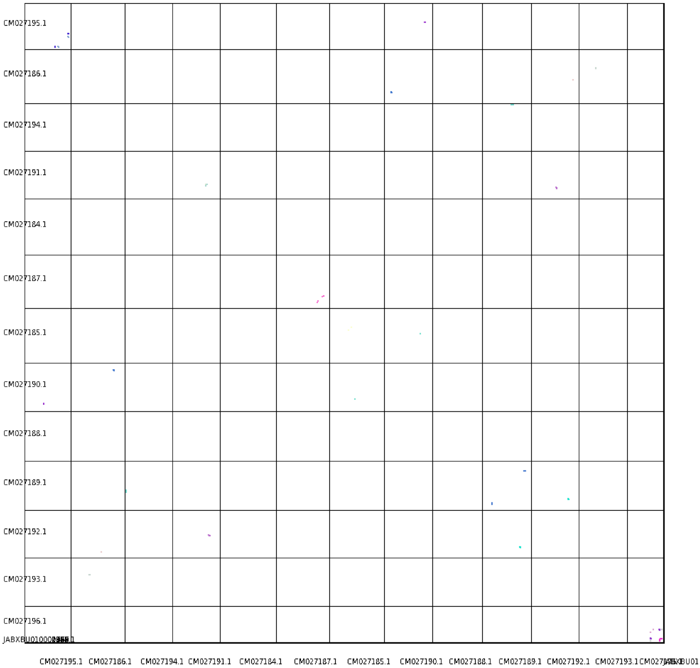

### *Bombyx mori*

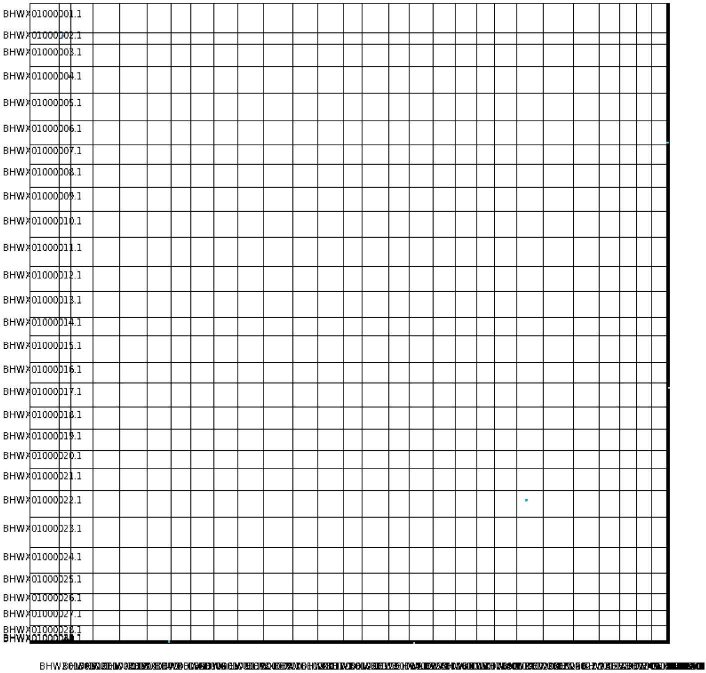

BHWX01000001.1
BHWX01000002.1
BHWX01000003.1
BHWX01000004.1
BHWX01000005.1
BHWX01000006.1
BHWX01000007.1
BHWX01000008.1
BHWX01000009.1
BHWX01000010.1
BHWX01000011.1
BHWX01000012.1
BHWX01000013.1
BHWX01000014.1
BHWX01000015.1
BHWX01000016.1
BHWX01000017.1
BHWX01000018.1
BHWX01000019.1
BHWX01000020.1
BHWX01000021.1
BHWX01000022.1
BHWX01000023.1
BHWX01000024.1
BHWX01000025.1
BHWX01000026.1
BHWX01000027.1
BHWX01000028.1

### *Carcinoscorpius rotundicauda*

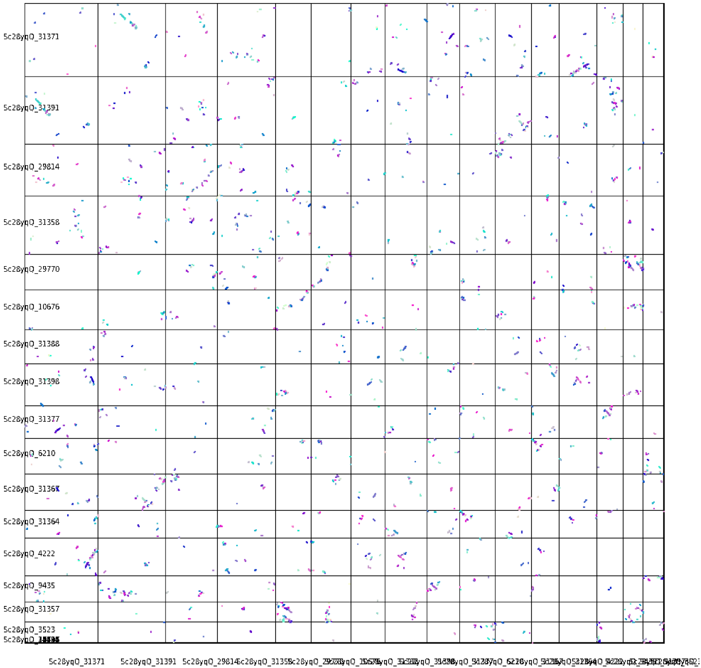

### *Centruroides sculpturatus*

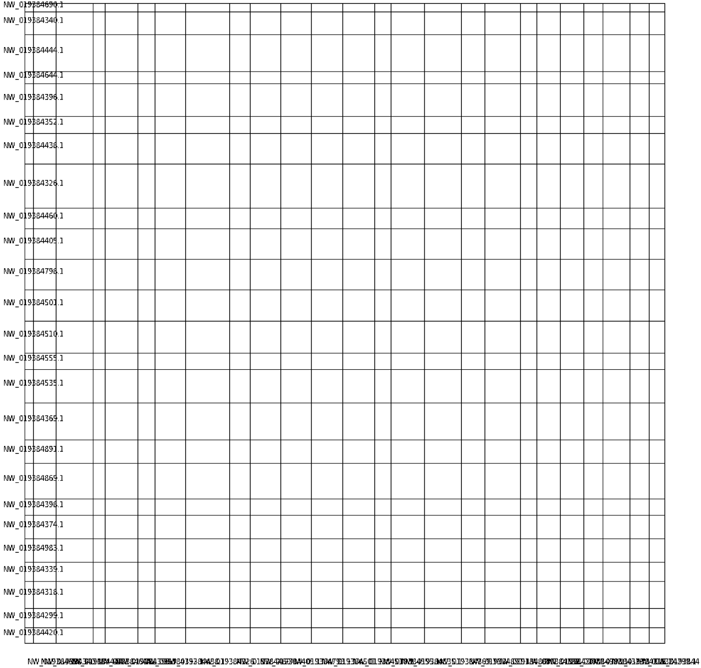

### *Drosophila melanogaster*

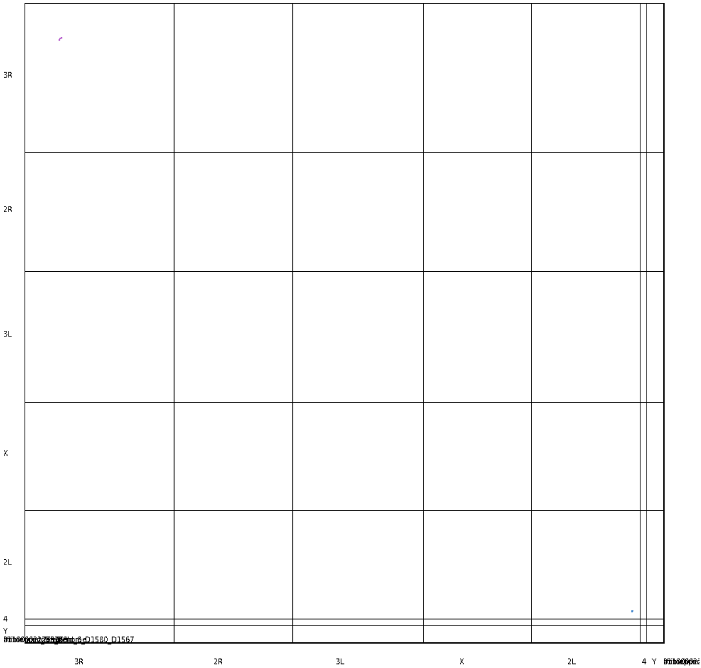

### *Haemaphysalis longicornis*

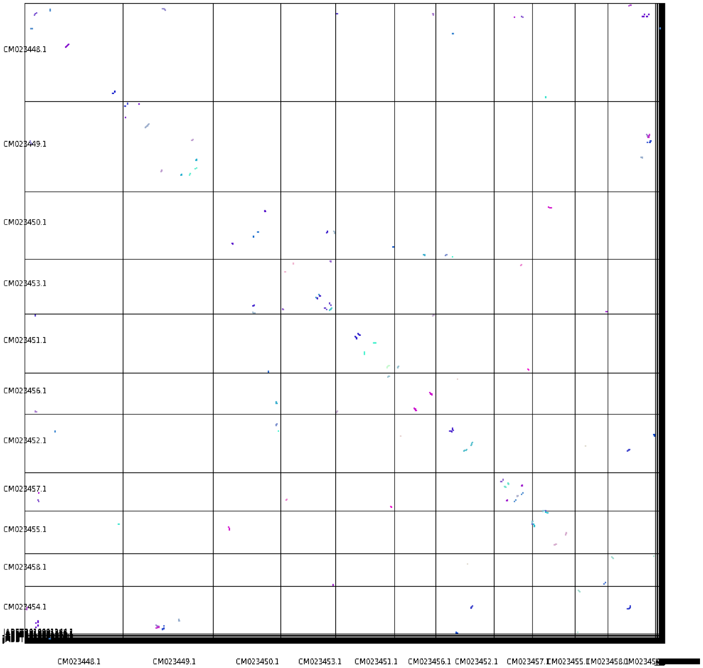

### *Ixodes scapularis*

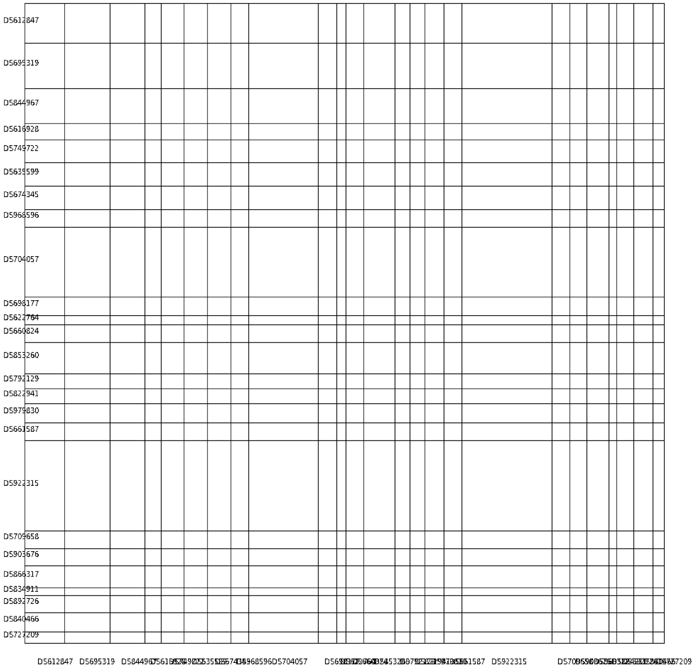

### *Latrodectus hesperus*

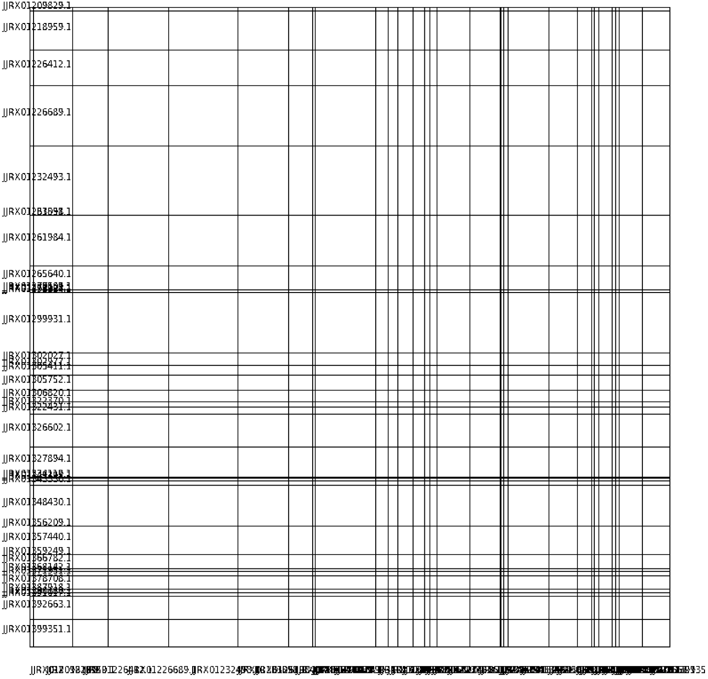

### *Limulus polyphemus*

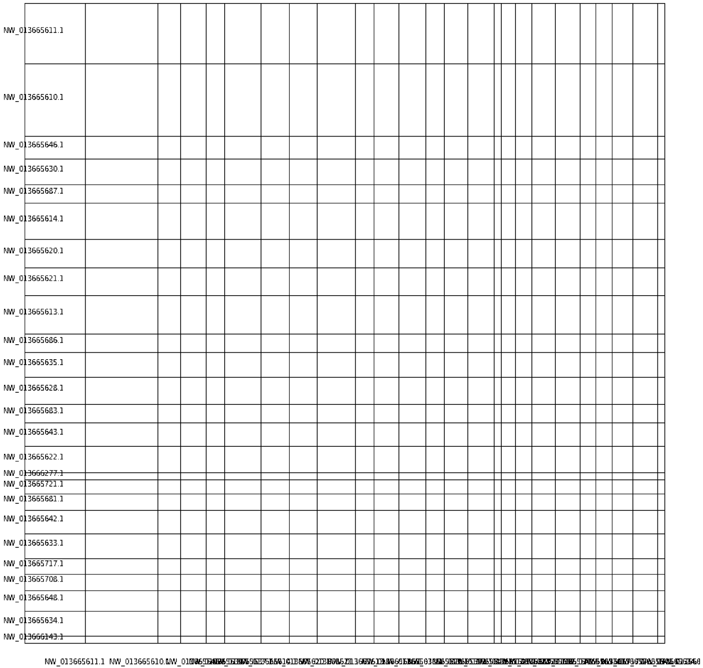

### *Loxosceles reclusa*

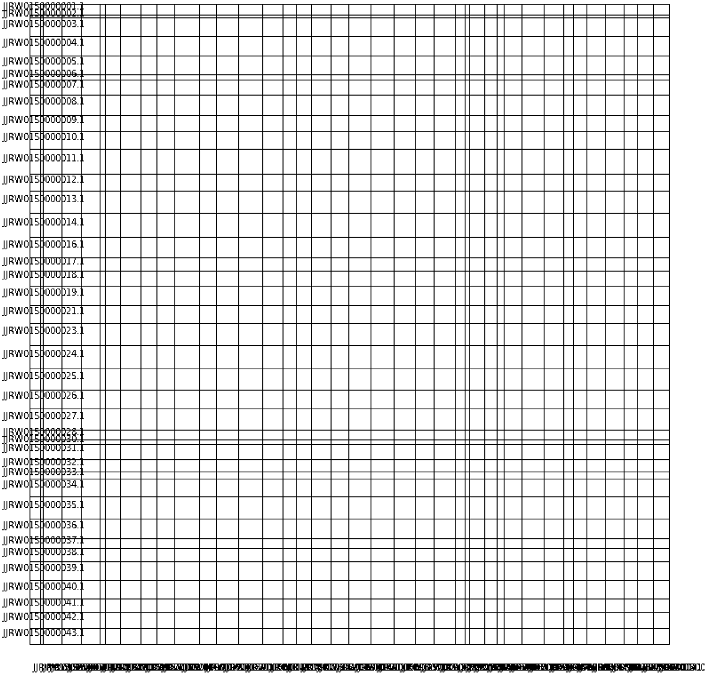

### *Metaseiulus occidentalis*

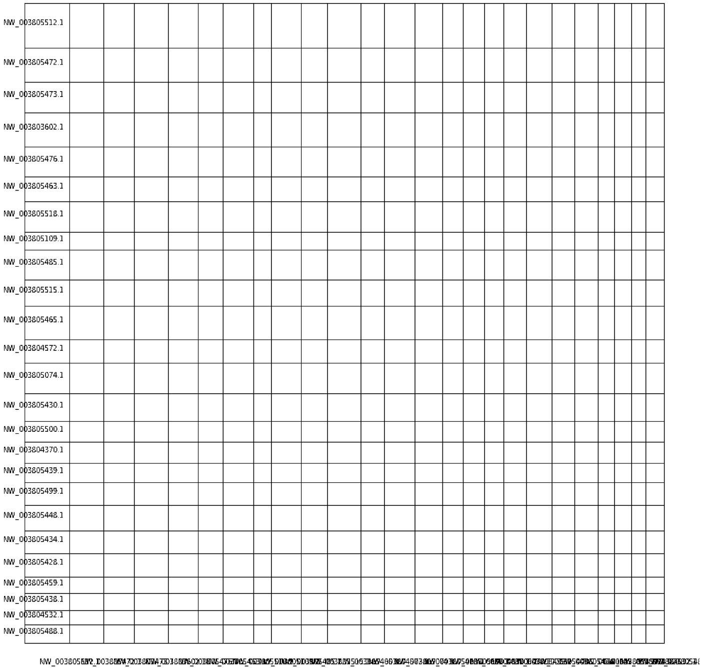

### *Nephila clavipes*

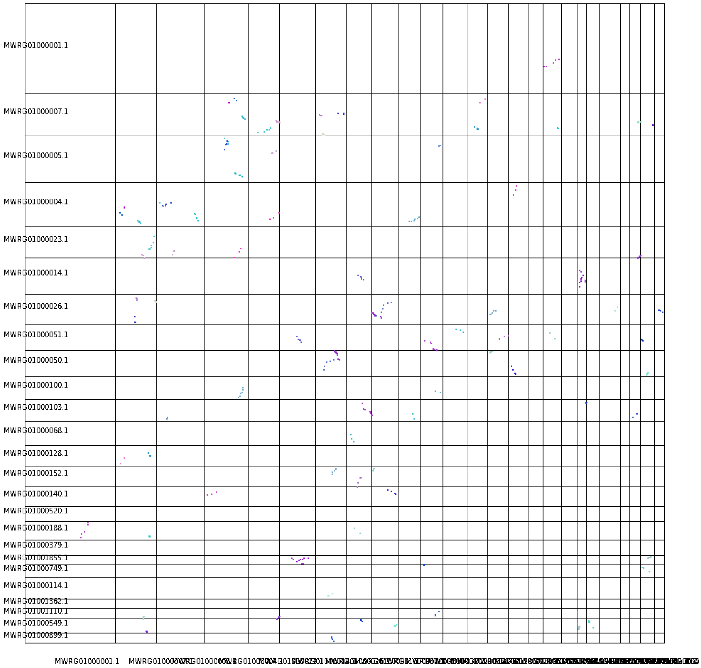

### *Parasteatoda tepidariorum*

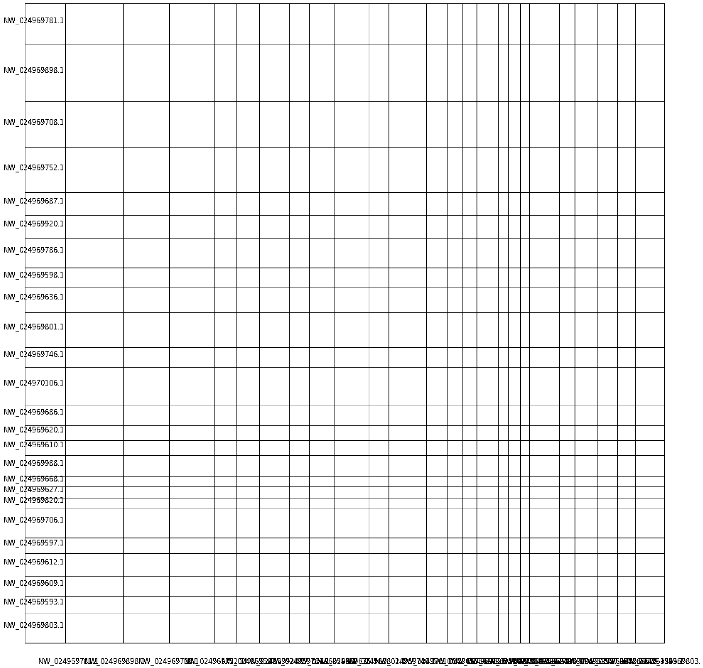

### *Stegodyphus mimosarum*

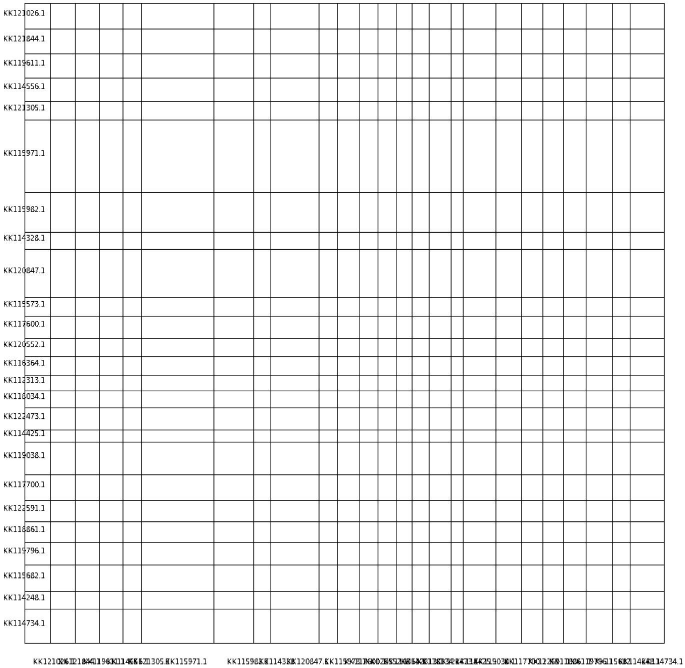

### *Sarcoptes scabiei*

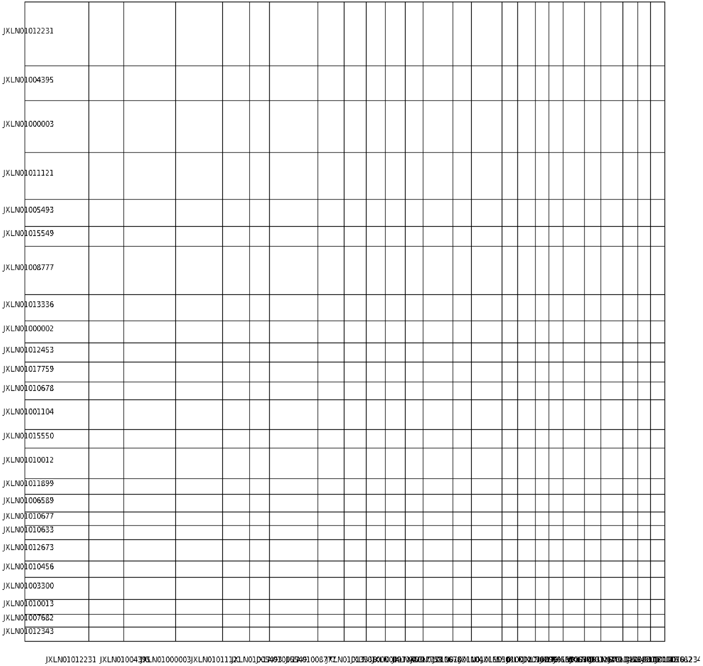

### Trichonephila antipodiana

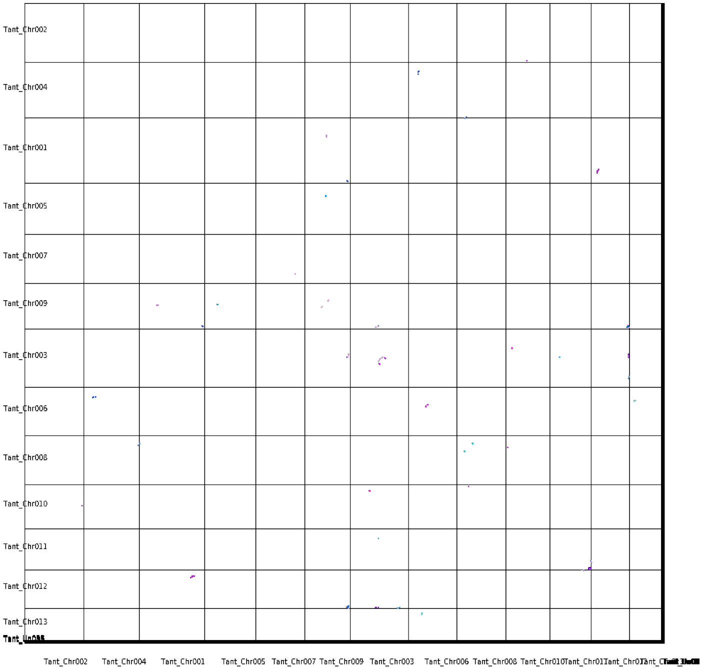

### *Tachypleus gigas*

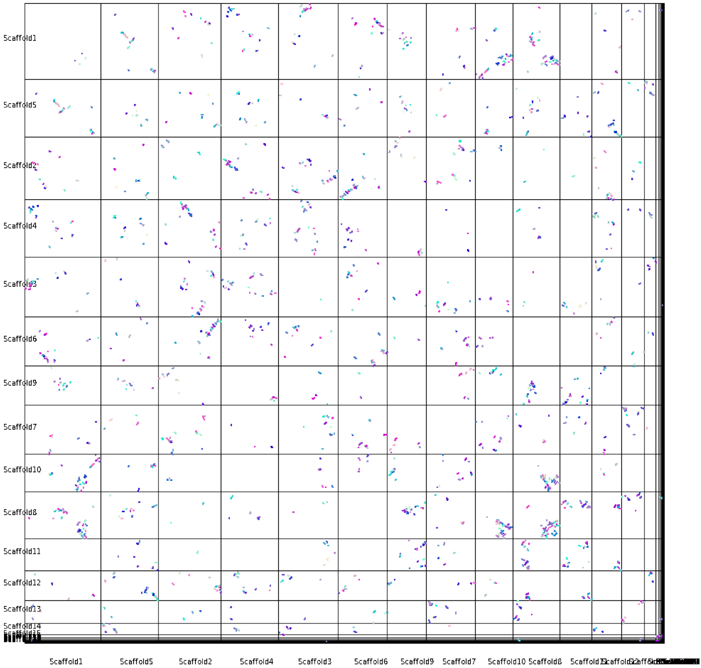

### Tetranychus urticae

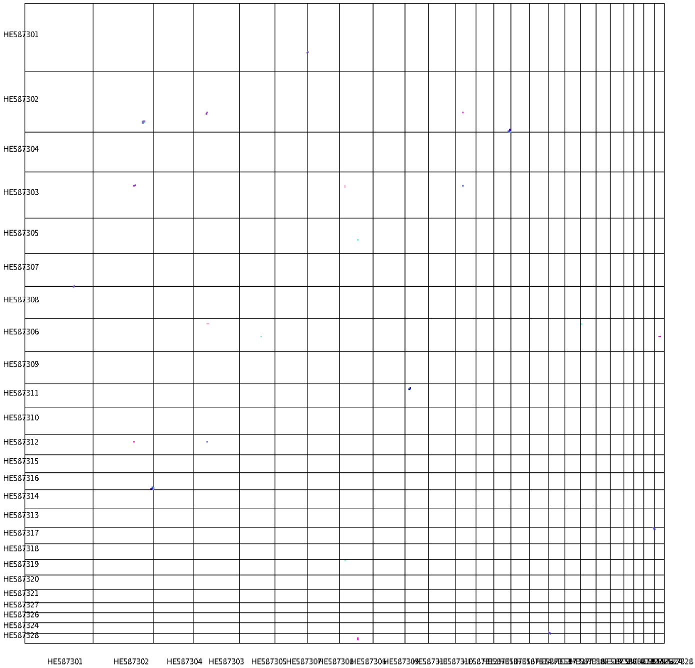

### *Varroa destructor*

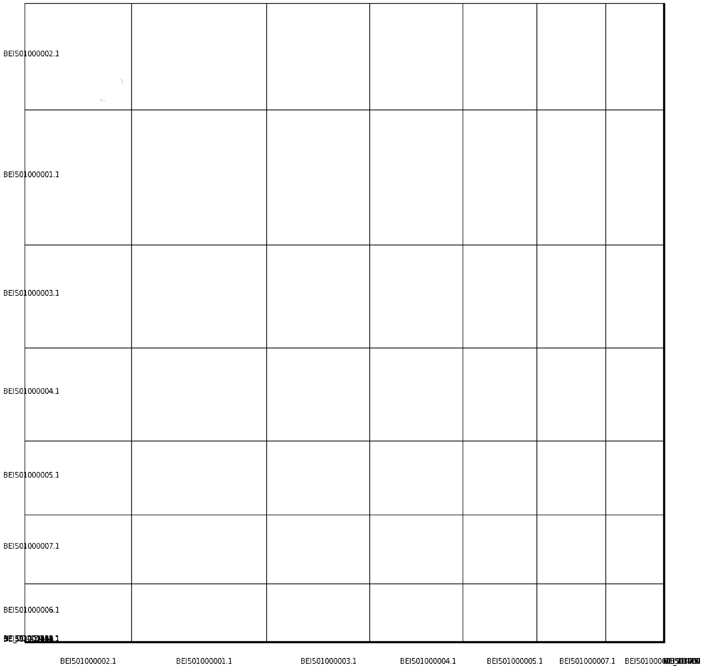
