## Supplemental Figure S6 for "A comprehensive examination of Chelicerate genomes reveals no evidence for a whole genome duplication among spiders and scorpions"

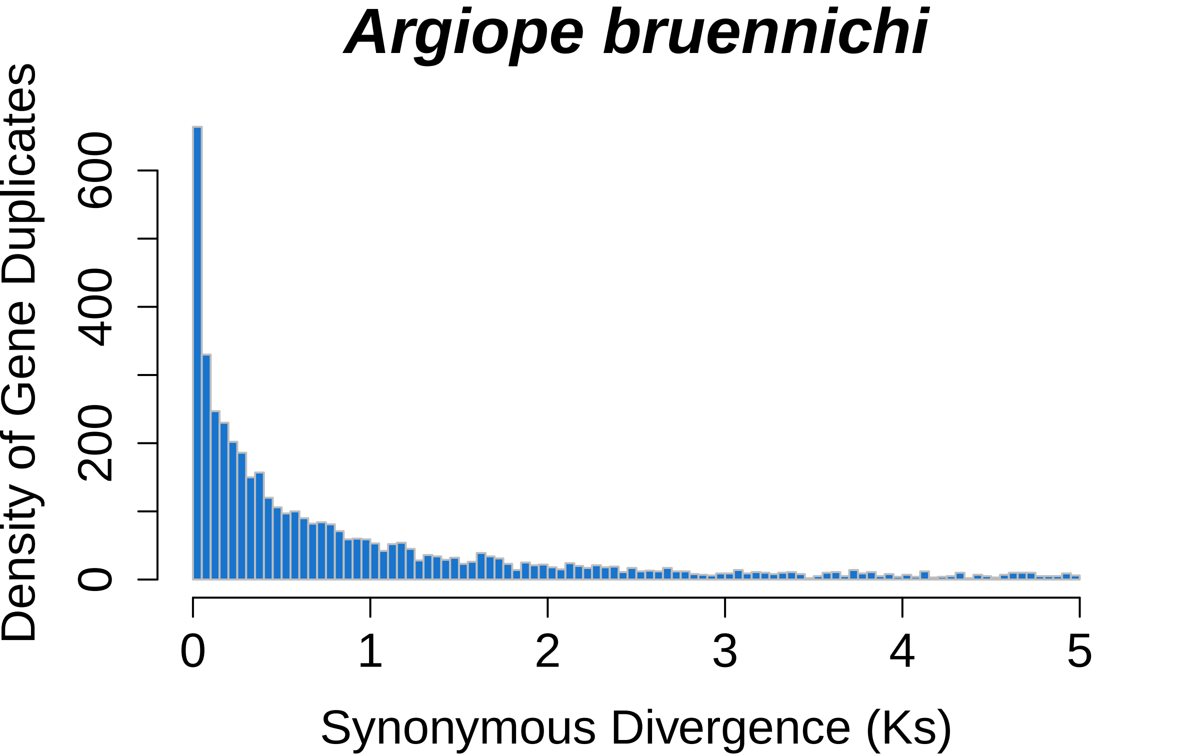
Argiope bruennichi
Density of Gene Duplicates
0
200
400
600
0
1
2
3
4
5
Synonymous Divergence (Ks)

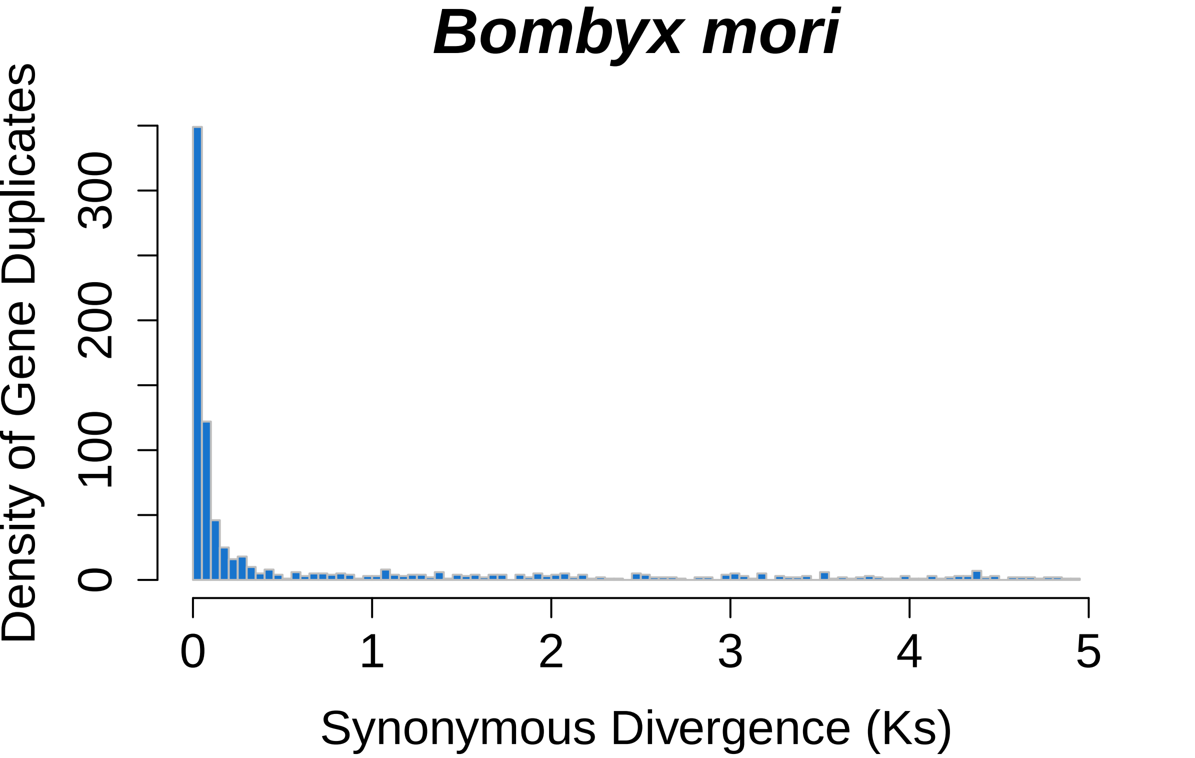

Bombyx mori
Density of Gene Duplicates
0
100
200
300
0
1
2
3
4
5
Synonymous Divergence (Ks)

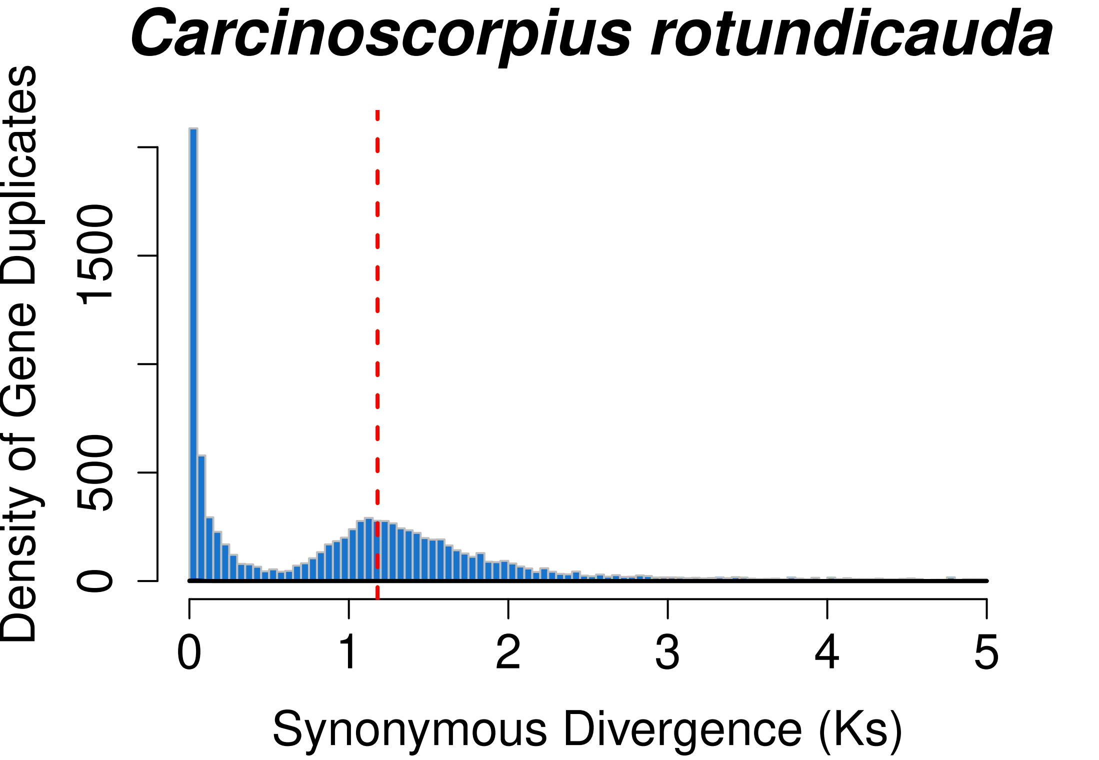

Carcinoscorpius rotundicauda
Density of Gene Duplicates
0
500
1500
0
1
2
3
4
5
Synonymous Divergence (Ks)

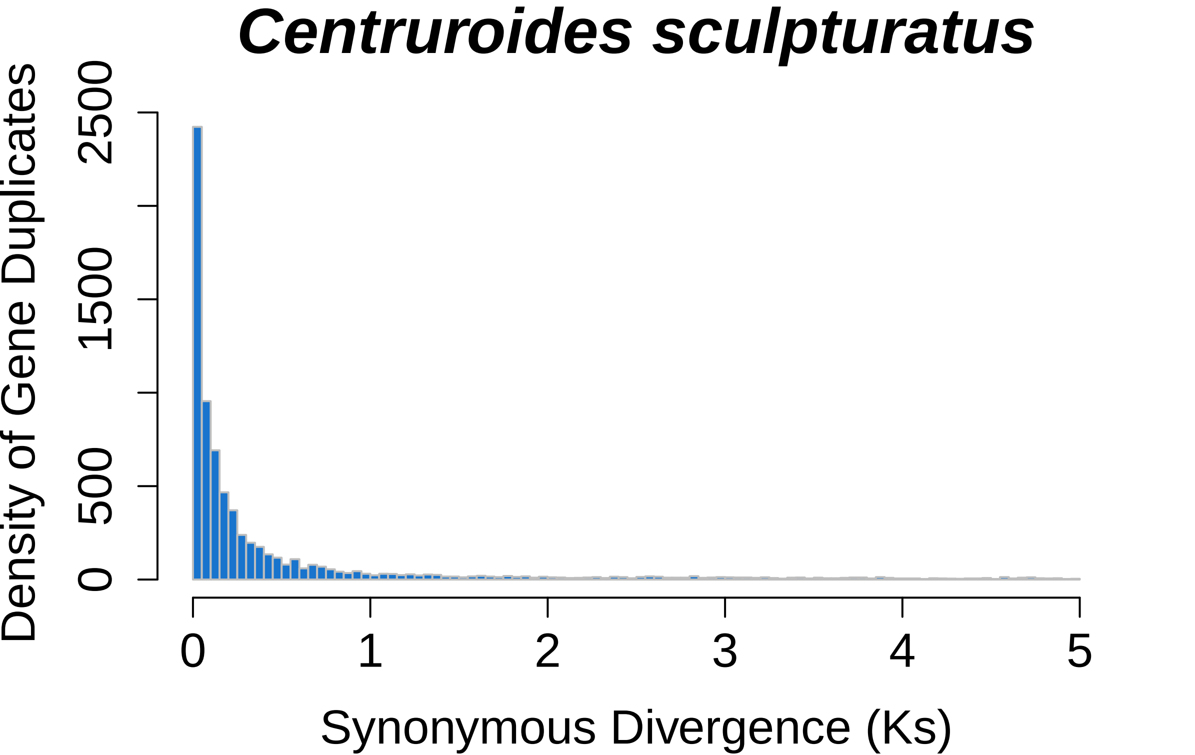

Centruroides sculpturatus
Density of Gene Duplicates
0
500
1500
2500
0
1
2
3
4
5
Synonymous Divergence (Ks)

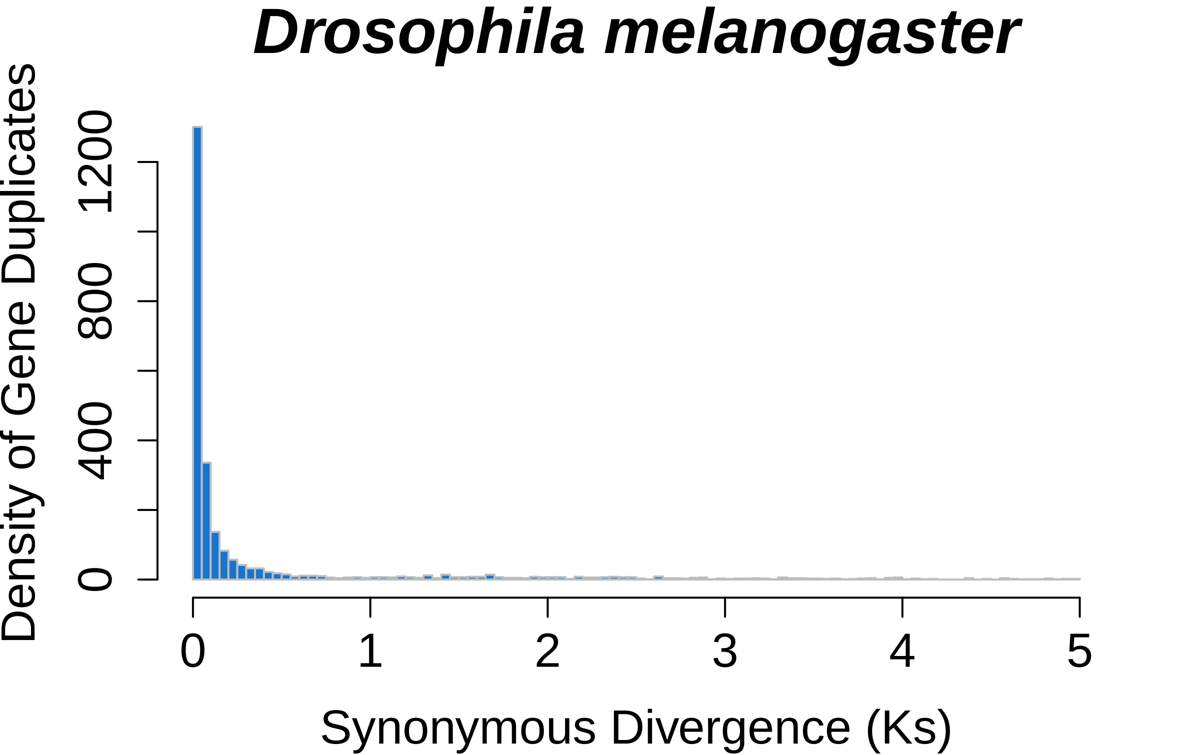
Drosophila melanogaster
Density of Gene Duplicates
0
400
800
1200
Synonymous Divergence (Ks)
0
1
2
3
4
5

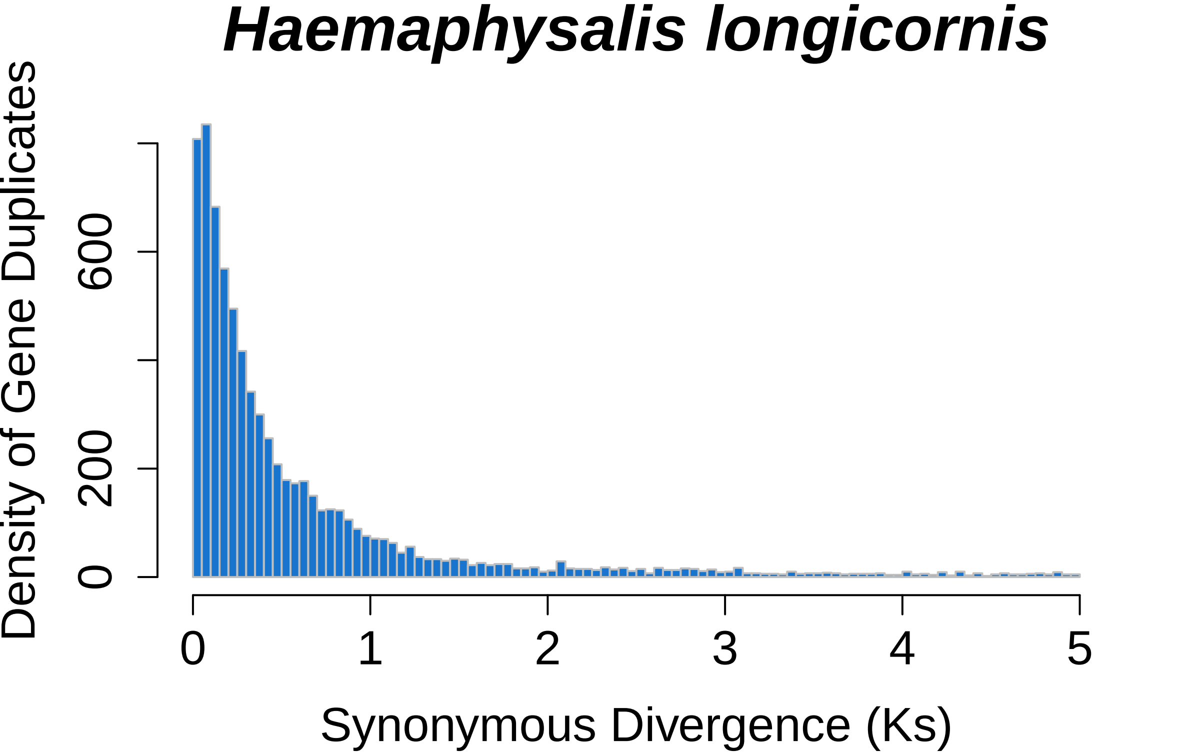

Haemaphysalis longicornis
Density of Gene Duplicates
0
200
600
0
1
2
3
4
5
Synonymous Divergence (Ks)

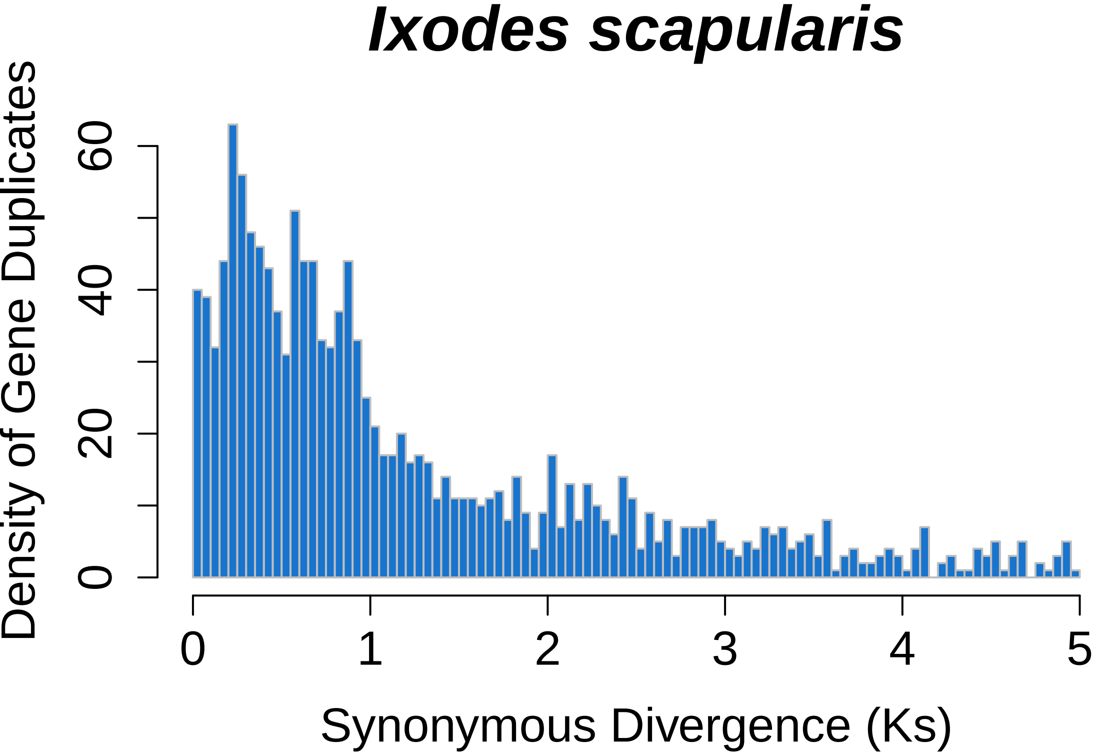

Ixodes scapularis
Density of Gene Duplicates
0
20
40
60
0
1
2
3
4
5
Synonymous Divergence (Ks)

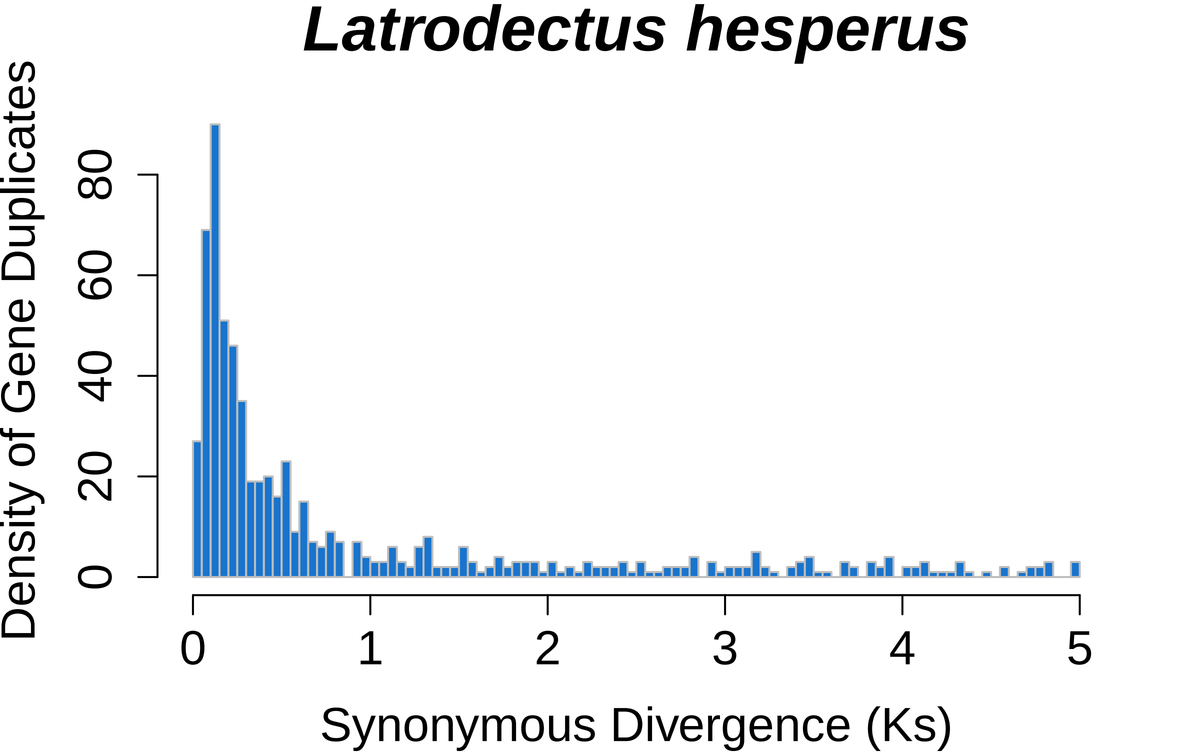
Latrodectus hesperus
Density of Gene Duplicates
0
20
40
60
80
0
1
2
3
4
5
Synonymous Divergence (Ks)

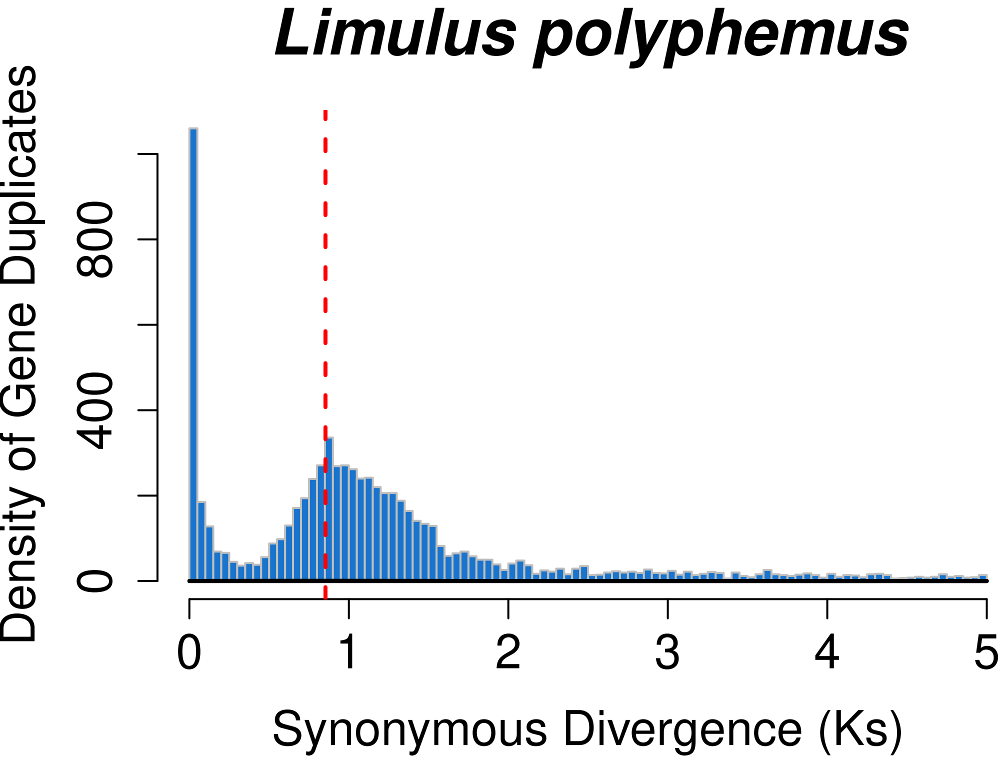

Limulus polyphemus
Density of Gene Duplicates
0
400
800
0
1
2
3
4
5
Synonymous Divergence (Ks)

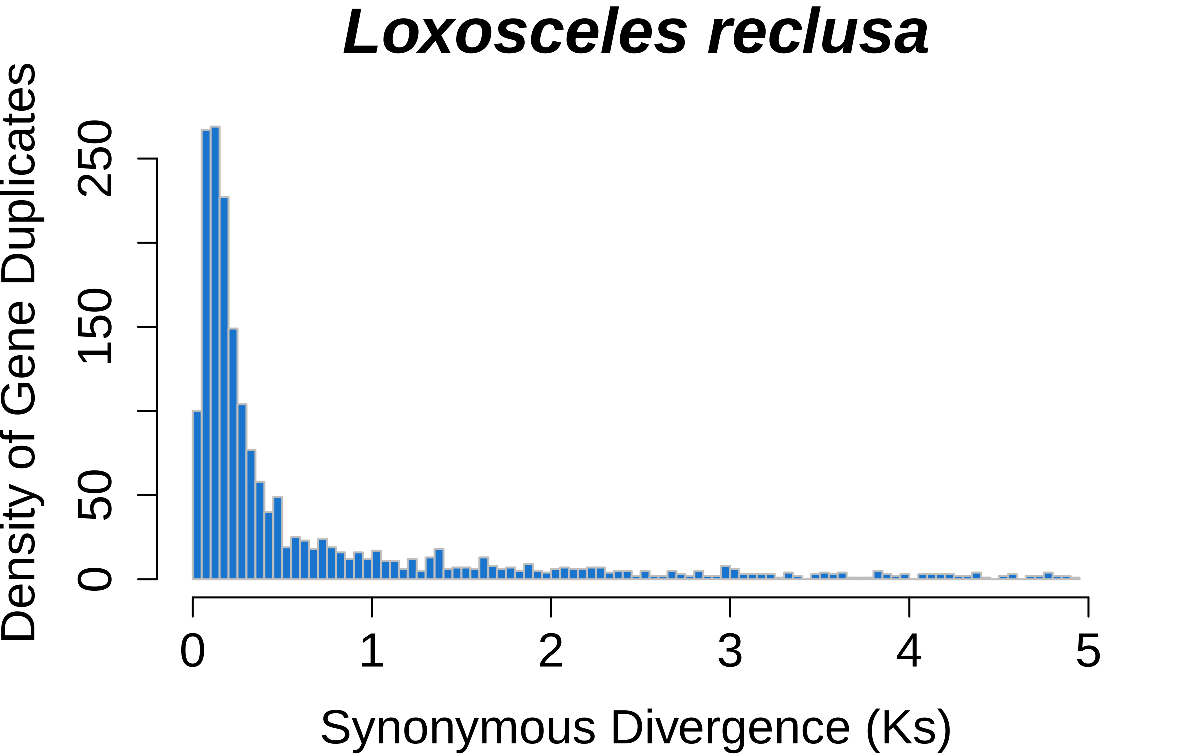

Loxosceles reclusa
Density of Gene Duplicates
0
50
150
250
0
1
2
3
4
5
Synonymous Divergence (Ks)

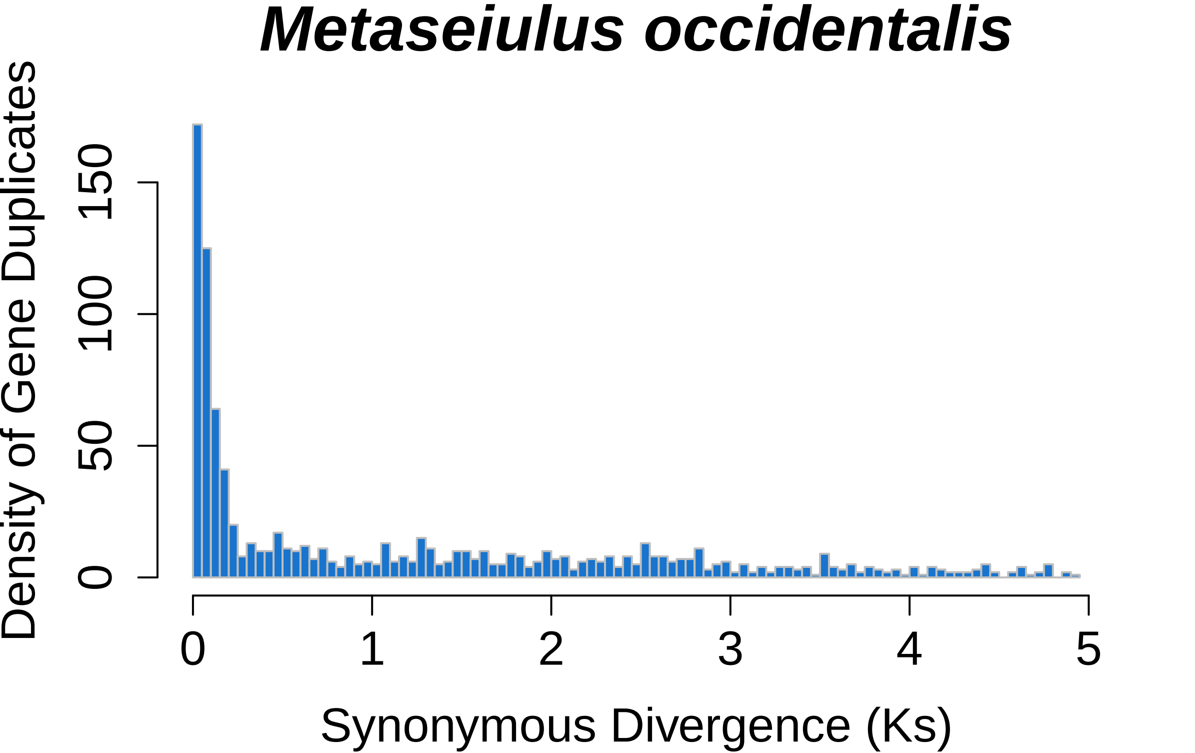

Metaseiulus occidentalis
Density of Gene Duplicates
0
50
100
150
0
1
2
3
4
5
Synonymous Divergence (Ks)

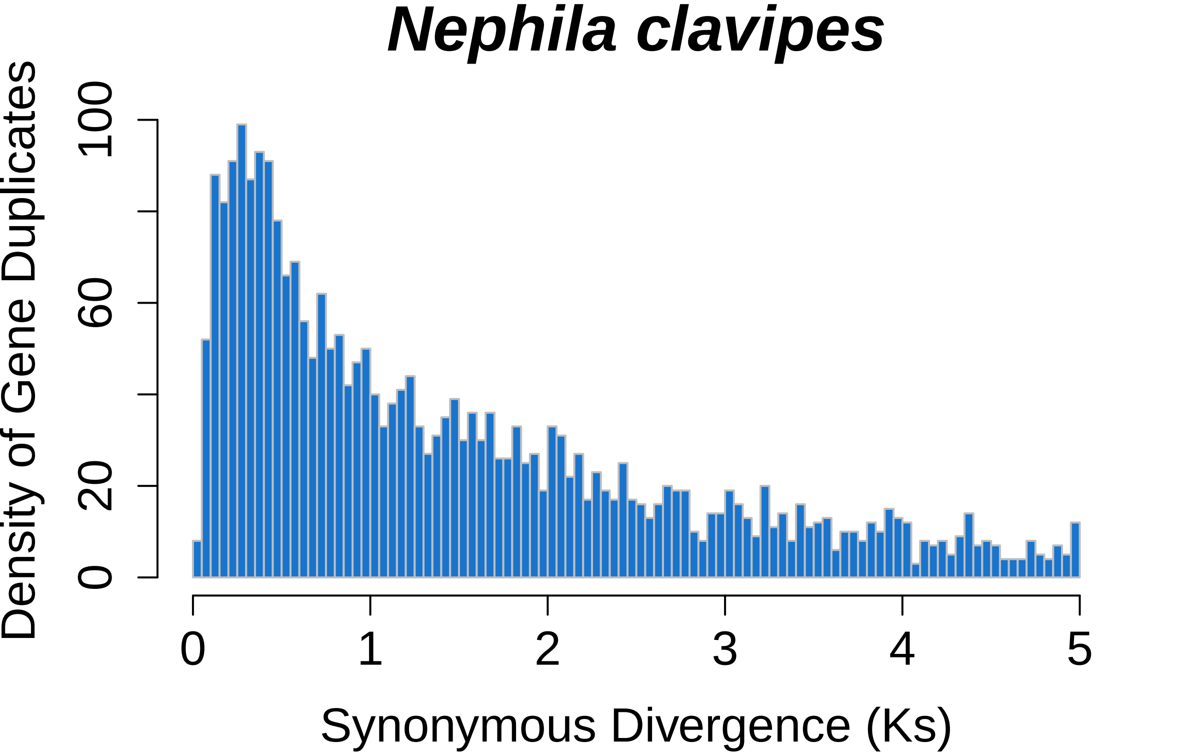
Nephila clavipes
Density of Gene Duplicates
0
20
60
100
0
1
2
3
4
5
Synonymous Divergence (Ks)

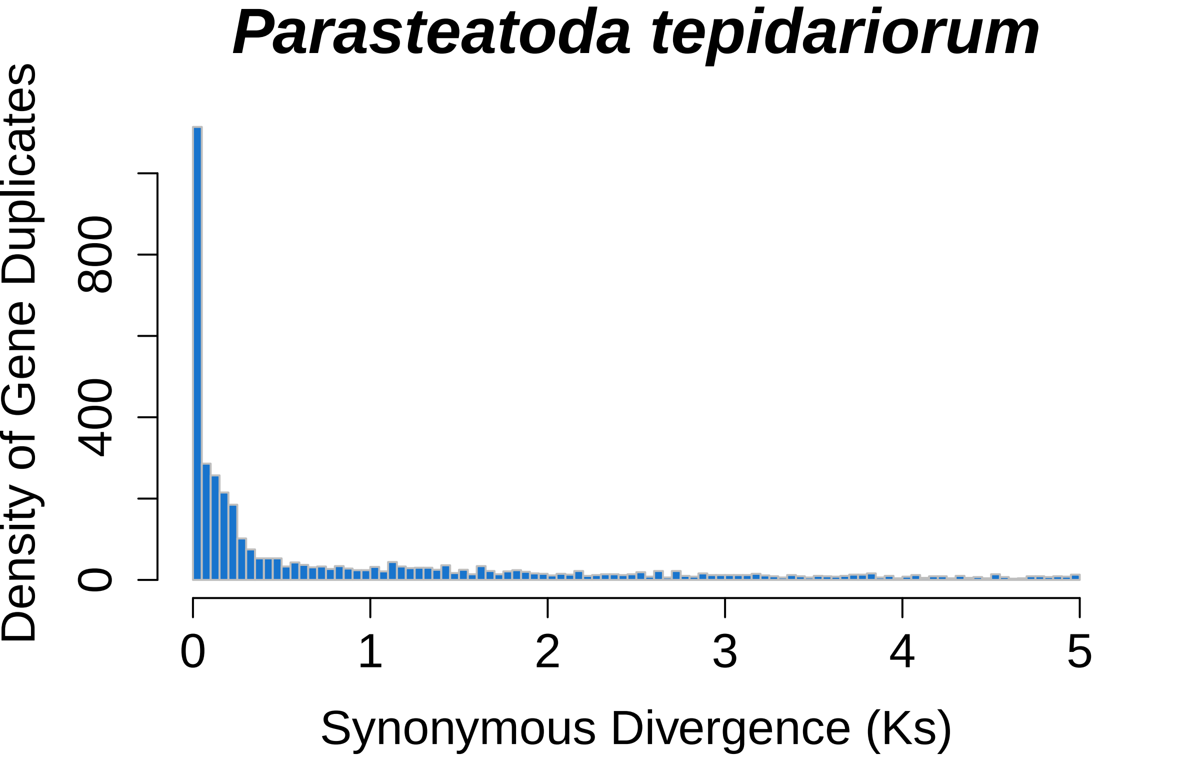
Parasteatoda tepidariorum
Density of Gene Duplicates
0
400
800
0
1
2
3
4
5
Synonymous Divergence (Ks)

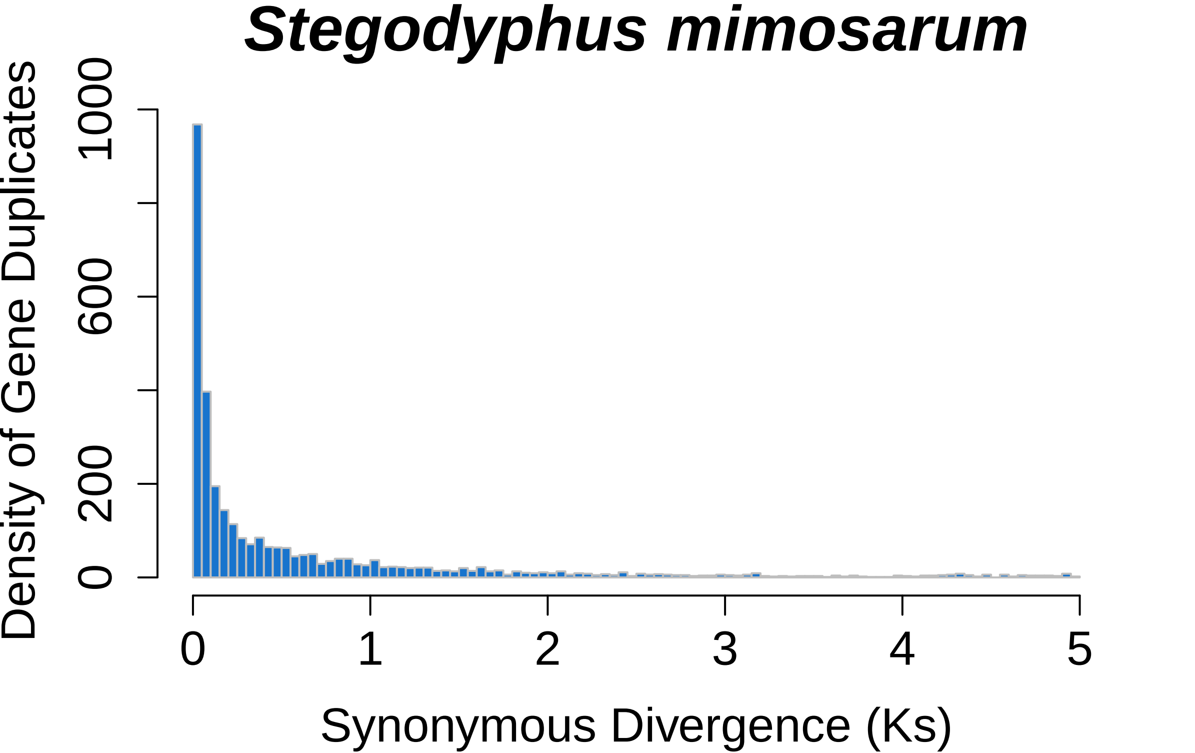

Stegodyphus mimosarum
Density of Gene Duplicates
0
200
600
1000
0
1
2
3
4
5
Synonymous Divergence (Ks)

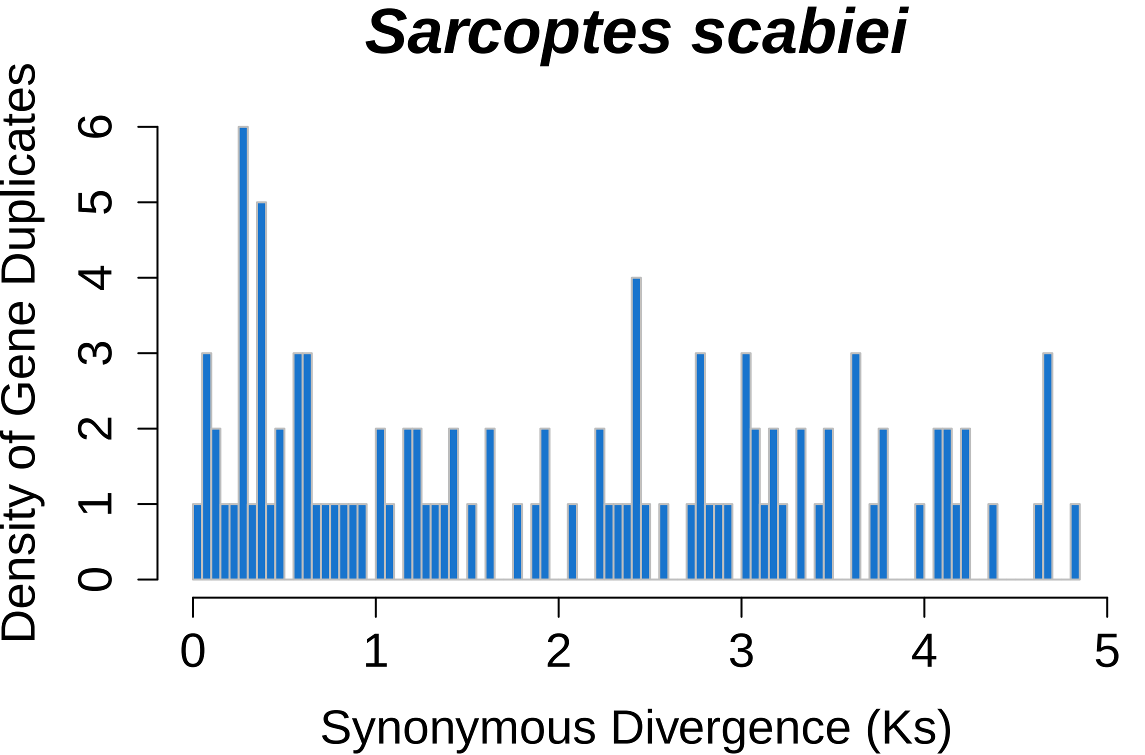

Sarcoptes scabiei
Density of Gene Duplicates
0
1
2
3
4
5
6
0
1
2
3
4
5
Synonymous Divergence (Ks)

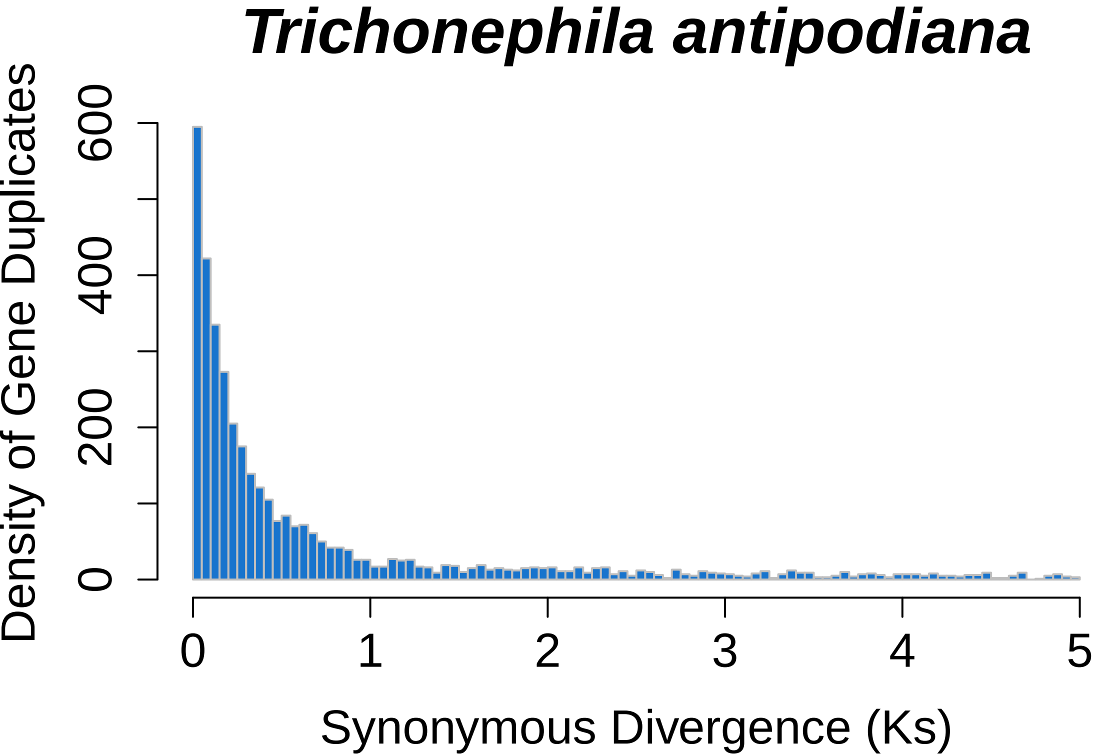

Trichonephila antipodiana
Density of Gene Duplicates
0
200
400
600
0
1
2
3
4
5
Synonymous Divergence (Ks)

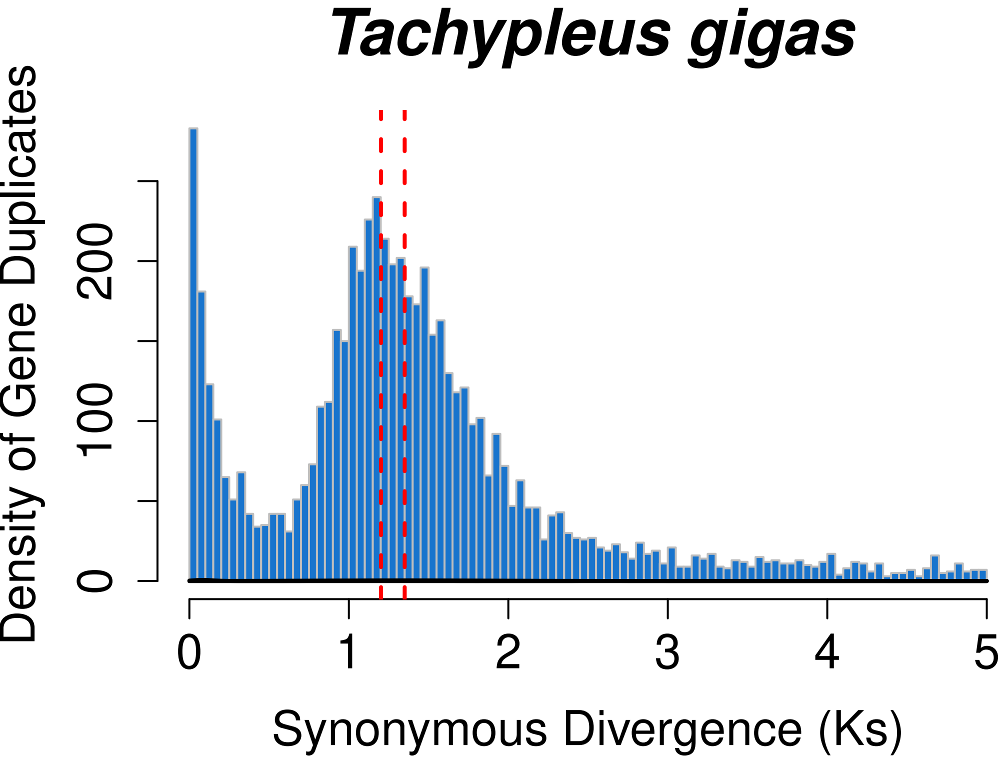
Tachypleus gigas
Density of Gene Duplicates
0
100
200
0
1
2
3
4
5
Synonymous Divergence (Ks)

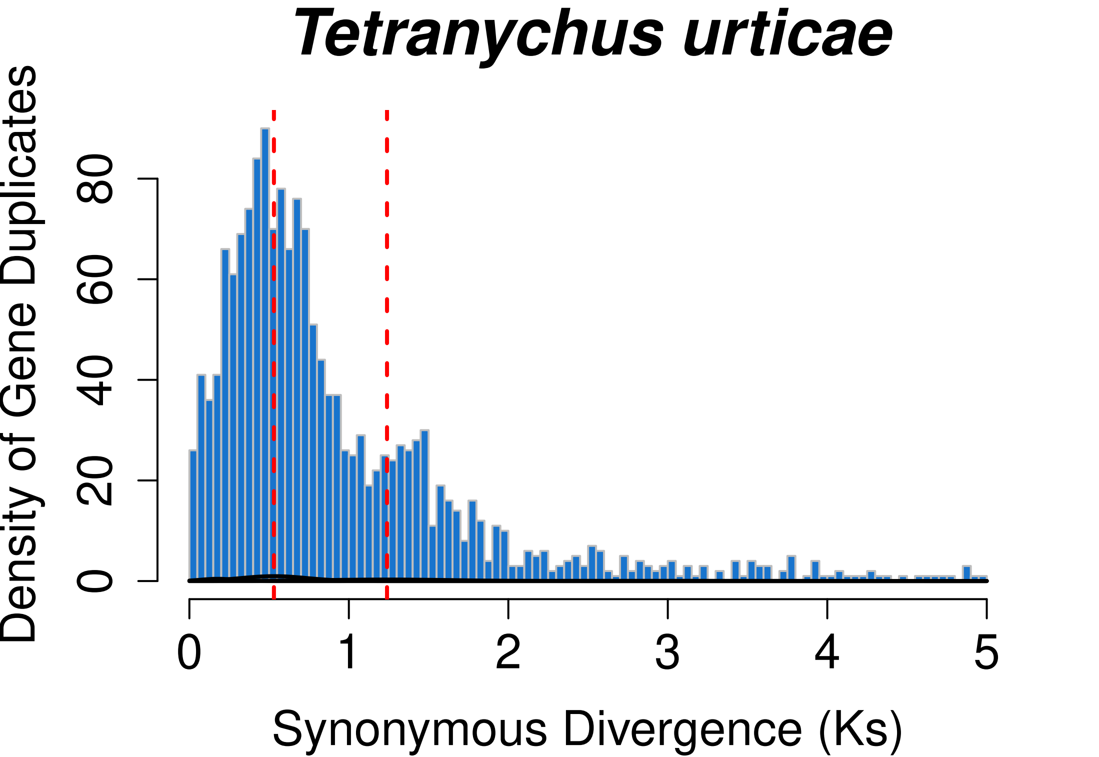
Tetranychus urticae
Density of Gene Duplicates
0
20
40
60
80
0
1
2
3
4
5
Synonymous Divergence (Ks)

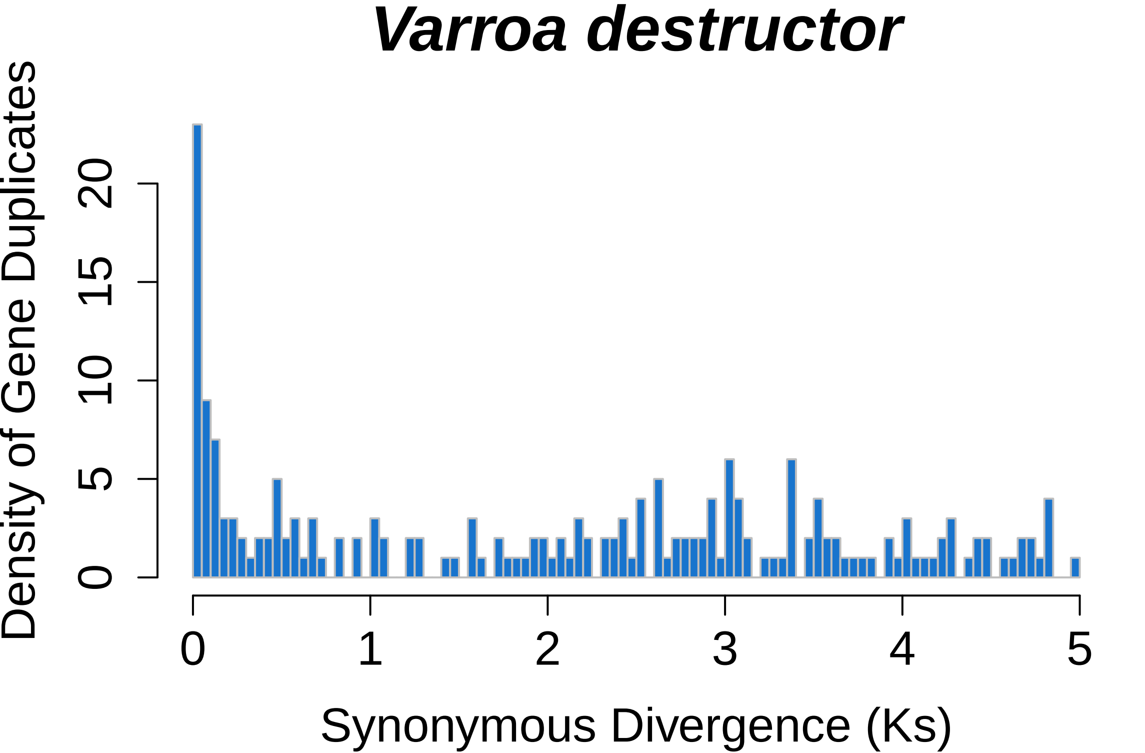
Varroa destructor
Density of Gene Duplicates
0
5
10
15
20
0
1
2
3
4
5
Synonymous Divergence (Ks)
